# Heliaphen, an outdoor high-throughput phenotyping platform designed to integrate genetics and crop modeling

**DOI:** 10.1101/362715

**Authors:** Florie Gosseau, Nicolas Blanchet, Didier Varès, Philippe Burger, Didier Campergue, Céline Colombety, Louise Gody, Jean-François Liévin, Brigitte Mangin, Gilles Tison, Patrick Vincourt, Pierre Casadebaig, Nicolas Langlade

## Abstract

Heliaphen is an outdoor pot platform designed for high-throughput phenotyping. It allows automated management of drought scenarios and plant monitoring during the whole plant cycle. A robot moving between plants growing in 15L pots monitors plant water status and phenotypes plant or leaf morphology, from which we can compute more complex traits such as the response of leaf expansion (LE) or plant transpiration (TR) to water deficit. Here, we illustrate the platform capabilities for sunflower on two practical cases: a genetic and genomics study for the response to drought of yield-related traits and a simulation study, where we use measured parameters as inputs for a crop simulation model. For the genetic study, classical measurements of thousand-kernel weight (TKW) were done on a sunflower bi-parental population under water stress and control conditions managed automatically. The association study using the TKW drought-response highlighted five genetic markers. A complementary transcriptomic experiment identified closeby candidate genes differentially expressed in the parental backgrounds in drought conditions. For the simulation study, we used the SUNFLO crop simulation model to assess the impact of two traits measured on the platform (LE and TR) on crop yield in a large population of environments. We conducted simulations in 42 contrasted locations across Europe and 21 years of climate data. We defined the pattern of abiotic stresses occurring at this continental scale and identified ideotypes (i.e. genotypes with specific traits values) that are more adapted to specific environment types. This study exemplifies how phenotyping platforms can help with the identification of the genetic architecture of complex response traits and the estimation of eco-physiological model parameters in order to define ideotypes adapted to different environmental conditions.

## Introduction

The use of fertilizers, irrigation and pesticides mitigated the effects of climatic hazards and had a large and positive impact in productivity gains between 1960 and 2000 (Foley et al., 2005; Tilman et al., 2002). Currently, because of the simultaneous need to reduce inputs in agricultural systems and the climatic uncertainty caused by climate change, the variability of farming conditions increased as compared to late century conditions. Both strategies of developing highly plastic crop genotypes or predicting the best combination of genotypes and agro-management practices to adapt to local conditions are thus key challenges. To tackle them and regardless of the magnitude of changes required in farming systems (Duru et al., 2015), characterizing and understanding how plants adapt to their environment is a necessary step.

In Europe, sunflower is mostly a rainfed crop and water deficit is frequently the main limiting factor (Blanchet et al., 1981; Debaeke et al., 2017). Climate change scenarios indicate that summer precipitation will decrease substantially in Southern and Central Europe and to a smaller degree in Northern Europe. However, during spring and autumn, precipitation change should be marginal. Overall, the intensity of daily precipitation should increase substantially (Moriondo et al., 2010). Drought episodes will thus continue to occur, probably with increasing variability from year to year.

In an agricultural context, a drought-tolerant plant is one that maintains crop production during gradual and moderate soil water deficits, most often without exhibiting protection mechanisms (Tardieu, 2011). Drought tolerance is the result of integrated processes taking place at different timescales and having long-term impact on leaf growth and transpiration (Tardieu et al., 2018). Water deficit affects a large spectrum of plant functions such as transpiration, photosynthesis, leaf and root growth, and reproductive development (Chaves et al., 2003) through underlying physiological processes (e.g. cell division, primary and secondary metabolism) (Tardieu et al., 2018).

In sunflower, responses to drought have long been studied at the physiological level and more recently at the molecular level. Different researchers have characterized its impact on leaf development, transpiration, photosynthesis and biomass allocation (Adiredjo, Navaud, Philippe Grieu, et al., 2014; Adiredjo, Navaud, Stephane Muños, et al., 2014; Andrianasolo, Casadebaig, et al., 2016; Andrianasolo, Champolivier, et al., 2016; Pereyra-Irujo et al., 2008; Velázquez et al., 2017). These studies have identified differences between genotypes for these traits resulting in improved water use efficiencies. The molecular bases of these processes start to be described in particular at the transcriptomic level. This allowed highlighting the role of osmotic potential maintenance both under controlled conditions and in the field (Rengel et al., 2012), the importance of reactive oxygen species (Ramu et al., 2016), of the signaling pathways of ABA (Rengel et al., 2012; Sarazin et al., 2017) ethylene (Manavella et al., 2006) and jasmonate (Andrade et al., 2017; Marchand, 2014). In a holistic approach, Marchand *et al*. have inferred a gene regulation network for drought based on hormonal signaling and shown its role in the evolution of wild sunflowers and modern breeding (Marchand, 2014). However, this eco-physiological and molecular knowledge is based on on simple drought stresses occurring mostly during early vegetative development. This is necessary to continue to develop it on more complex and realistic scenarios, taking into account the dynamic and organ-level natures of drought stresses and plant responses.

Over the population of cropping environments, the different soils and climates combined with chosen variety and management conditions generate highly diverse water deficit scenarios (e.g. Chenu et al., 2013). A given trait can therefore have a positive, null or even negative impact on crop performance depending on drought scenarios (Casadebaig, Zheng, et al., 2016). Because plant traits of interest are context dependent, we need tools to measure and integrate the multiple and overlapping mechanisms involved in plant response to water deficit, among which plant growth and transpiration are the primary targets (Tardieu et al., 2018). To address this objective, phenotyping platforms and crop mathematical model are complementary tools.

Driven by technological development, phenotyping platforms are tools created to efficiently measure plant traits while controlling cultivation conditions. Greenhouse-based platforms allow a very fine control of plant environment (light, water, nutriments) (Cobb et al., 2013; Granier et al., 2005; Pieruschka and Poorter, 2012) and were successfully used to conduct association genetics studies (Cabrera-Bosquet et al., 2012) and derive plant trait ranges that can serve for crop modeling (Lenz-Wiedemann et al., 2010; Steduto et al., 2009). However, design options that are important for achieving a high-throughput (small plots and plants) and a sound control of environmental conditions (closed facility, artificial lighting) drives cultivation conditions away from field-conditions. This gap may cause difficulties when generalizing plant response to agricultural conditions (Mittler, 2006).

Computer-based plant modeling approaches have recently emerged as a method to complement and improve the resource-limited experimental exploration of the adaptation landscape (Chapman et al., 2003; Hammer et al., 2006; Messina et al., 2006, 2011). These software models are based on mathematical equations that represent the biological processes linked to plant growth and development as a function of time, environment (climate, soil, and management), and genotype-dependent parameters. Those parameters are expected to be more heritable than complex traits, which are more subjected to environmental variation and to genotype-environment interaction (Heslot et al., 2014). Accordingly, simulation can be used as a tool to predict trait × trait and trait × environment interactions and help to assess traits value when scaling up from the individual plant to the plant population (one field) and finally up to the population of fields in the cropping area. Simulation successfully allowed to cluster ressource availability time-series in stress scenarios at a continental scale (Chenu et al., 2013), to search for adapted traits for specific targeted environment types (Casadebaig, Mestries, et al., 2016; Chapman et al., 2002), and to infer trait values beyond field experiments and available genetic diversity (Casadebaig, Zheng, et al., 2016; Martre et al., 2015).

In this context, we aim to characterize various sunflower genotypes using a phenotyping platform to measure genotype-dependent traits, thus enabling the simulation of genotypes performance for a diversity of abiotic stress scenarios. Ultimately, linking phenotyping, crop modeling and simulation would allow us to design a decision support system to better target genotypes to their most performing cropping environment, accounting for climatic uncertainty.

To achieve this goal, we developed the Heliaphen phenotyping platform to measure key phenotypic trait involved in plant response to water deficit for large panels of genotypes. In order to study traits related to crop performance (e.g. seed number and mass, seed oil content), the plants are grown in outdoor condition and we rely on automation to manage water deficit scenarios and measure plant traits. Daily measurements of leaf and plant growth rate, plant transpiration rate and water deficit level allowed to estimate phenotypic response to water deficit (reaction norms). For example, we measured how leaf expansion and plant transpiration varied with water deficit (Casadebaig et al., 2008), estimated the parameters of such response curves using non-linear regression and used these genotype-dependent parameters in a crop simulation model (SUNFLO, Casadebaig et al., 2011) to simulate the performance of phenotyped genotypes in non-observed environments.

Here, we validated our ability to manage distinct drought scenarios impacting plant leaf area, biomass and yield related traits (green box in figure 1). Two distinct approaches were then undertaken as case studies to demonstrate the platform capacities. The first approach concerns genetics and genomics, where we used an association study and transcriptomics to identify genetic basis of the response of yield-related traits to water deficit (blue box in figure 1). The second one is a crop modeling and simulation study to illustrate how traits measured using the phenotyping platform can be used in a crop simulation model to analyse how they impact yield on diverse and non-observed cropping conditions (pink box in figure 1).

**Figure 1.**
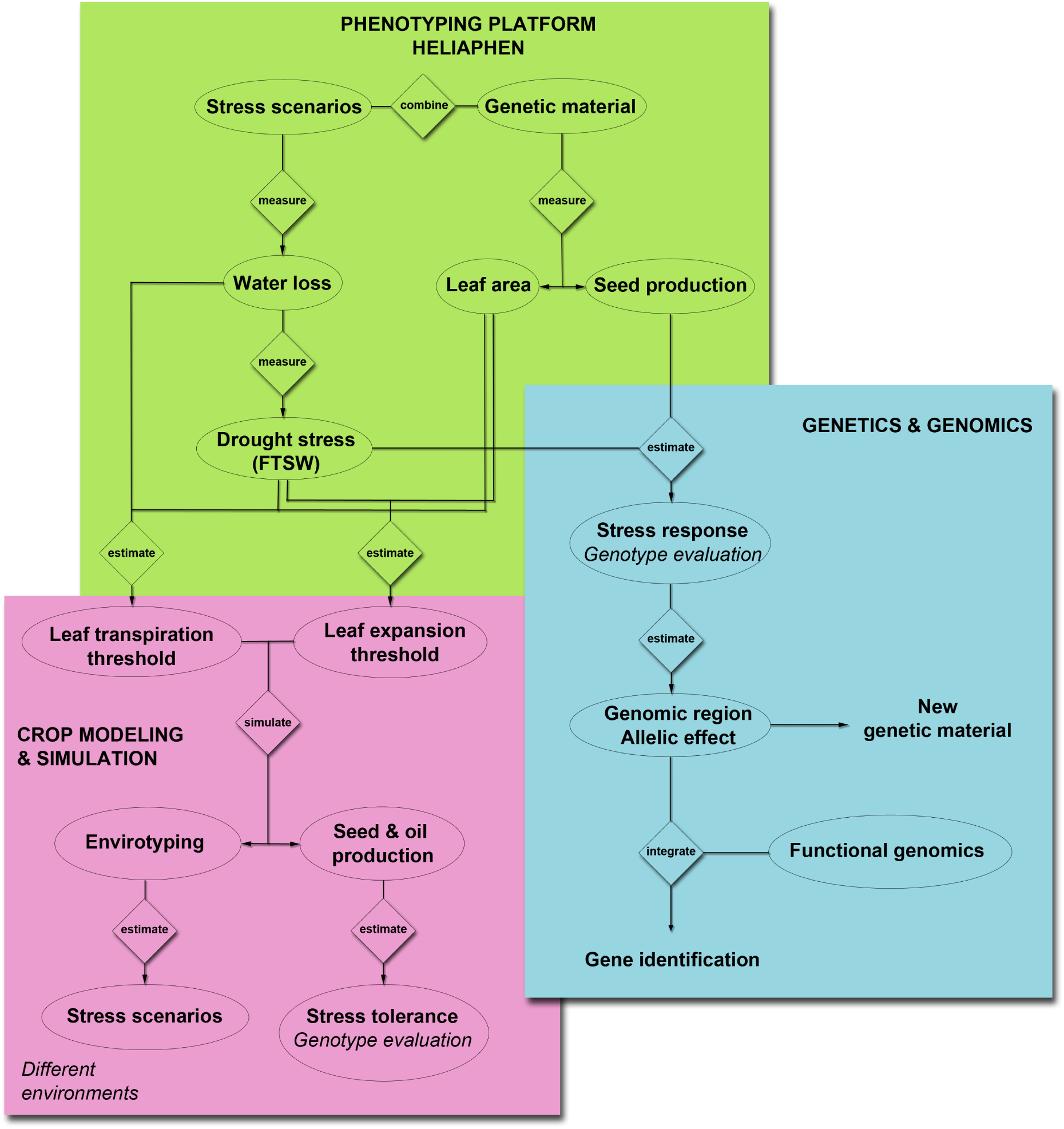
Overview of the approaches targeted with the Heliaphen phenotyping platform. This figure illustrates how we used the phenotyping platform to (1) grow diverse genetic material under designed stress scenarios and measure key ecophysiological traits and plant water status (green box), (2) conduct genetic and genomics studies on measured phenotypes (blue box) to identify gene functions, and (3) estimate crop simulation parameters based on the measured phenotypes to be used in simulation studies. Simulation allows to characterize abiotic stress at the crop level (envirotyping) and evaluate genotype value (performance and stress tolerance) in diverse cropping conditions (pink box).

## Materials & Methods

### Heliaphen Platform

Heliaphen is a 650 m^2^ outdoor phenotyping platform on which a robot performs measurements and irrigates plants grown in pots. Heliaphen can hold up to 1300 plants spread over 13 blocks consisting of 2 rows of 50 pots of 15 liters (figure 2A). This outdoor platform is surrounded by nets to break wind and prevent bird entry. In order to control the water balance and manage water deficit scenarios, pots are covered with a cone shaped cap to prevent rainwater from entering the pot. Environmental conditions are recorded and logged (temperature, wind, precipitation, evaporative demand).

**Figure 2.**
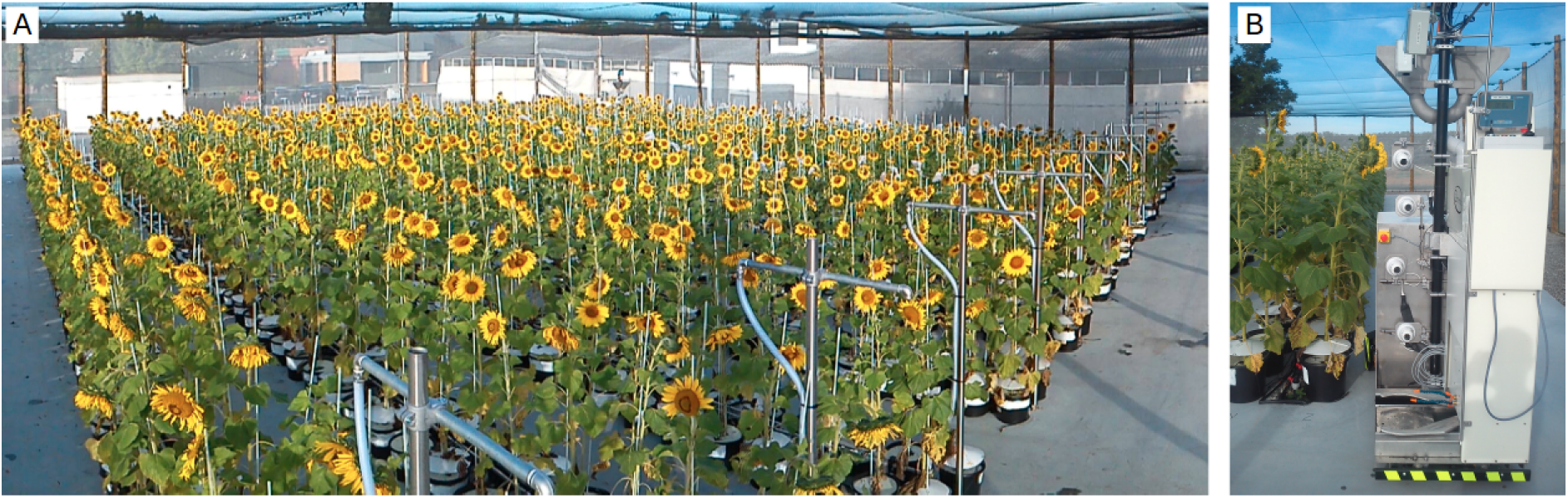
General view of the Heliaphen platform. (A) General view of the Heliaphen platform with an area of 650 m2 and 1300 plants spread over 13 blocks consisting of 2 rows of 50 pots. (B) The Heliaphen robot, able to grab pots, weigh them, water and phenotype individual plants.

The robot (figure 2B) is based on an electrical propulsion system with batteries allowing about 48h of continuous functioning before recharging (about 4h). It is equipped with a 60 liters water tank used to manage irrigation. Navigation on the platform, water and electricity management are fully autonomous. The robot is equipped with a clamp to grab pots, a digital scale (Midrics 1, Sartorius Weighting Technology GmbH, Gettingen, Germany) and four digital cameras (Prosilica GC650 with Sony ICX424 sensor and 659×493 resolution). Weighing, irrigating and image acquisition for one plant lasts between 30 and 90 seconds depending on the amount of water to apply. The weight of the pots is recorded before and after irrigation, along with plant pictures from the cameras.

The robot manages drought scenarios at the plant level, i.e. for each pot. Before the beginning of the stress period (early vegetative stage), pots are irrigated to saturation, and after excessive water is drained, pots are weighed to estimate pot mass at full soil water capacity. For each plant, the planned water deficit level, expressed in fraction of transpirable soil water (FTSW, Sinclair et al. (2005)) is converted to a target pot weight and logged into the planification software. For each interaction of the robot with the plants, the robot navigate in the plant stand, grabs a pot with its clamp and weighs it, then irrigate the plant according to the defined water deficit scenario, as the difference between the actual and the target pot weight. Both control and stressed plants are processed one to four times per day by the robot to estimate plant transpiration according to the experiment size and required phenotyping intensity. In addition, an independent dripping irrigation system is used to allow plant growth before robot-dependent, controlled irrigation management. It allows different irrigation and fertilization regimes using liquid nutrient supply with a Dosatron system.

Beside to the high-throughput automated measurements, manual measurements are currently involved for other plant traits, like individual leaf area and plant height. The platform was developed using sunflower as a model species but maize, soybean and tomato have been successfully tested on the Heliaphen platform.

### Plant material, experiments and multi-environment trials

Various panels of sunflower plant material and different experiments have been used in the different part of this study, their main characteristics are summarized in Table 1 and shortly described below, with the full description available in Badouin et al. (2017) supplementary material. Each trial has an identifier consisting of two numbers encoding the year, followed by two letters encoding the location (HP for experiment on the Heliaphen platform, EX and RV for field experiments), the last two numbers are the trial number.

**Table.1.**
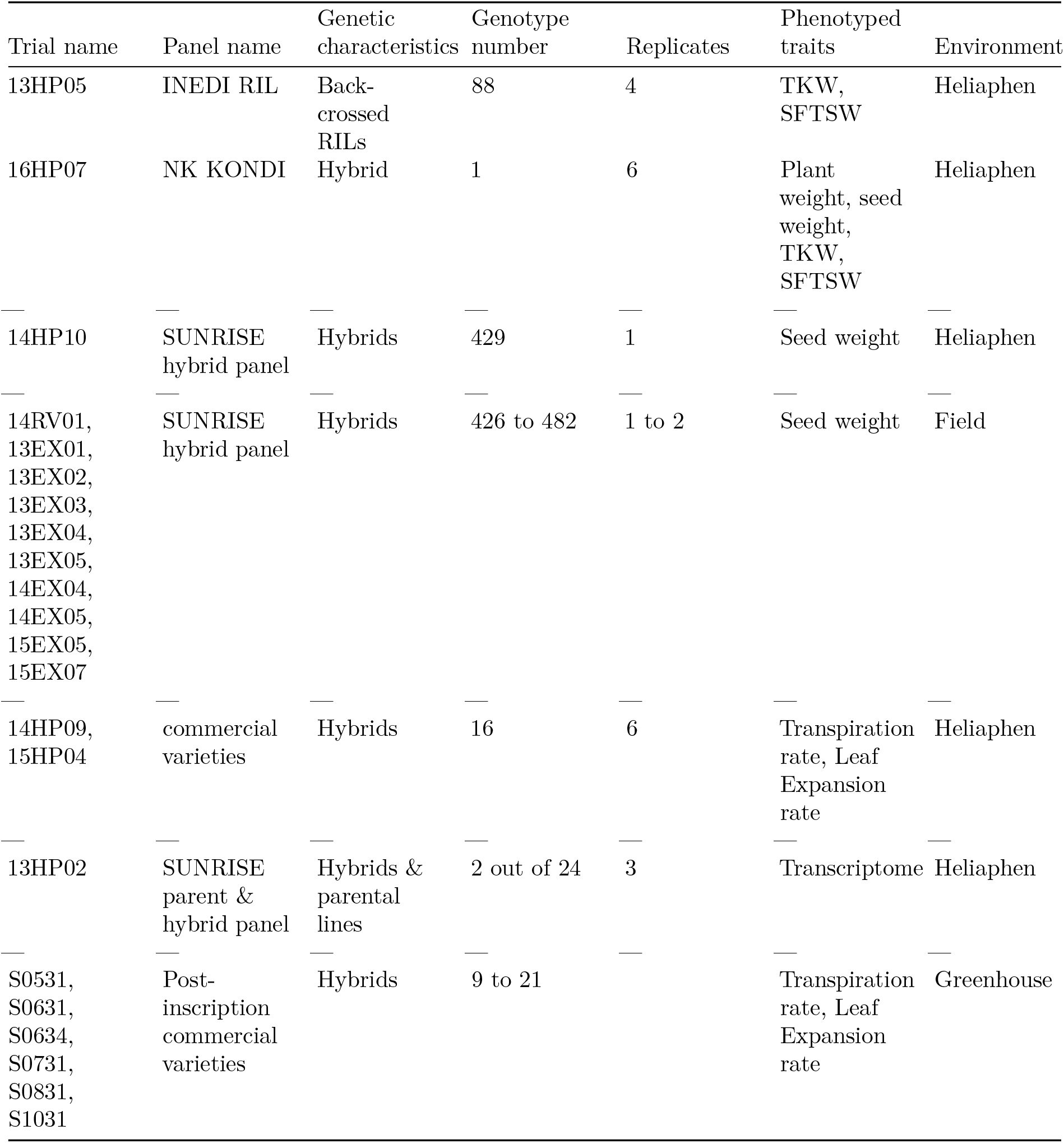
Plant material and experiments used in this study.

| Trial name | Panel name | Genetic characteristics | Genotype number | Replicates | Phenotyped traits | Environment |
| --- | --- | --- | --- | --- | --- | --- |
| 13HP05 | INEDI RIL | Back-crossed RILs | 88 | 4 | TKW, SFTSW | Heliaphen |
| 16HP07 | NK KONDI | Hybrid | 1 | 6 | Plant weight, seed weight, TKW, SFTSW | Heliaphen |
| 14HP10 | SUNRISE hybrid panel | Hybrids | 429 | 1 | Seed weight | Heliaphen |
| 14RV01, 13EX01, 13EX02, 13EX03, 13EX04, 13EX05, 14EX04, 14EX05, 15EX05, 15EX07 | SUNRISE hybrid panel | Hybrids | 426 to 482 | 1 to 2 | Seed weight | Field |
| 14HP09, 15HP04 | commercial varieties | Hybrids | 16 | 6 | Transpiration rate, Leaf Expansion rate | Heliaphen |
| 13HP02 | SUNRISE parent & hybrid panel | Hybrids & parental lines | 2 out of 24 | 3 | Transcriptome | Heliaphen |
| S0531, S0631, S0634, S0731, S0831, S1031 | Post-inscription commercial varieties | Hybrids | 9 to 21 |  | Transpiration rate, Leaf Expansion rate | Greenhouse |

#### General validation of the phenotyping platform

The comparison between Heliaphen platform and field experimentations for performance-related traits is based on trials 14HP10 and 14RV01 respectively. These two experiments were conducted on the same genetic panel (490 hybrids, table 1), sown at the same time (April 30 and May 5, 2014) and in the same geographical location (straight line distance between trials was 4.11 km). Others field experiments (13EX01 to 15EX07) were conducted on the same hybrid panel at different locations and sown on different dates. On the 14HP10 trial plant seed weight was estimated by sampling individual plants in each plots, whereas in other trials it was computed as the ratio of harvested seed weight to the number of plants per plot.

The platform capacity to generate distinct water deficit levels and their impact on performance related traits is based on trial 16HP07. This trial is based on one genotype, the commercial hybrid *NK KONDI*, subjected to different drought scenarios. We defined seven different levels of water stress applied during the reproductive stage (flowering - harvest) and six replicates for each stress. Target stress levels were 0.2, 0.3, 0.4, 0.6, 0.75, 0.9 and 1, expressed in FTSW with 1 corresponding to maximum plant available water (the plant is not water stressed).

#### Trials used for genetics and genomics approach

The genetic study was performed on the 13HP05 Heliaphen trial. The tested panel is a RIL population, coming from the cross between XRQ/B (SF193) and PSC8/R (SF326). To overcome disease sensitivity issues due to the duration of experiment and the sensitivity of PSC8 to Verticillium and Alternaria, and avoid branched plants, 84 RILs carrying the Rf1 fertility restoration allele were back-crossed to the non-branched male-sterile parent XRQ/A (SF193). We conducted two drought scenarios, the control and a constant water deficit level (FTSW = 0.4; 1) from flowering to harvest, and four replicates.

The transcriptomic study was performed on the 13HP02 Heliaphen trial. This trial was composed of 24 genotypes, including 4 females lines, 4 males lines and their 16 hybrids. The panel included the hybrid SF193xSF326 and its parental lines, on which we present results here. Two drought scenarios were defined during the vegetative stage, the control (irrigated) and a progressive water deficit (non-irrigated), with three replicates. The treatments started 35 days after germination, when irrigation was stopped for stressed plants. To ensure a comparable state of individual plant regarding water stress for transcriptomic analysis, pairs of stressed and control plants were harvested when the FTSW of the stressed plant reached 0.1.

#### Trials used for crop modeling and simulation

The comparison between Heliaphen platform and greenhouse experiments regarding response traits is based on trials 14HP09 and 15HP04 for the Heliaphen platform and S0531, S0631, S0634, S0731, S0831, S1031 for the greenhouse trials. In these trials, a total of 82 sunflower commercial hybrids were phenotyped for the response of leaf expansion and plant transpiration to water deficit. Because only three control genotypes were common among all trials, the comparison was performed on mean and variance of the population. To estimate the response traits for one genotype, we phenotyped the leaf area expansion rate, the transpiration rate and the water deficit level for twelve plants, growing under two water treatments. The two drought scenarios were defined during the vegetative stage, the control (irrigated) and a progressive water deficit (non-irrigated), with six replicates. The response traits is the regression of the phenotype (expansion and transpiration) response (ratio of each stressed plants on the average of control plants) on the water deficit level; the protocol is fully described in Casadebaig et al. (2008).

### Genetic association study

To demonstrate the capacity of identifying genetic markers associated to the water stress response of a yield component, we performed a genetic study on the 13HP05 experiment. The observation of the Thousand Kernel Weight (TKW) on the Heliaphen platform together with available genotyping data on this population (from Badouin et al., 2017) allowed us to perform a genome wide association study (GWAS). On the 13HP05 trial, the population of RILs was submitted or not to drought stress, the FTSW was maintained to 0.4 or 1 thanks to the robot during the seed filling stage. To study the TKW stress response, the model below is used to identify associated genetic markers:

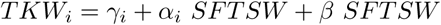

where *TKW_i_* is the thousand kernel weight for the *i*-th genotype, *SFTSW* is a stress variable (integration of 1 minus *FTSW*) observed, *γ_i_* is the genotypic effect associated to *i*-th genotype plus the residual error, *α_i_* is the coefficient of stress response associated at the i^th^ genotype, and *β* coefficient corresponds to the average response of *TKW* to *SFTSW*.

Association tests are done on a set of 2,240 independent markers corresponding to the individual genetic bins discriminated by the 84 studied RILs. These markers come from SNPs detected by genomic re-sequencing and implemented on an Affymetrix AXIOM genotyping array. All individuals have been genotyped using this array composed of 586,986 markers and results were presented for genome assembly in (Badouin et al., 2017). We conserved 55,951 markers which are homozygous in SF193 and heterozygous in SF326xSF193 and with homozygous marker frequency between 40 and 60 % and with less than 10% of heterozygocity or missing data. We obtained 2,240 merged markers assigned to 17 Linkage-groups thanks to Carthagene (Givry et al., 2004) and positioned on the sunflower genetic map (Badouin et al., 2017).

The association tests were performed using a multi-locus mixed-model, MLMM, proposed by Segura et al. (2012). The model is based on a classic GWAS model (Yu et al., 2006) on which marker selection is performed using a forward stepwise approach. At each step the SNP with the smallest p-value is added to the model and the p-value thus as the residual variance are re-estimated for all cofactors. The forward selection analysis stops when the proportion of variance explained by the model is close to zero. The analysis has been conducted thanks to the MLMM code written by Segura et al. (2012) and with ASReml-R package (Butler et al., 2009) and the modifications provided by (Bonnafous et al. 2017). The method used is now available on CRAN with *mlmm.gwas* package.

### Transcriptomics

A transcriptomic analysis was conducted on the 13HP02 trial to identify candidate genes underlying previously identified QTLs. We focused on the transcriptome of the SF193xSF326 hybrid and the SF193 parental line under two conditions, irrigated or non-irrigated as these genetic and stress conditions correspond to the studied situation in the 13HP05 genetic study. The transcriptomes of the hybrid was compared with the parent line subjected to stress conditions. Transcriptome analysis was done on plant pairs (stress-control) when the FTSW of the stressed individual reaches 0.1.

Leaves were harvested between 11:00 and 13:00. To select the leaf to harvest, the total leaf number was estimated, and leaves at the rank position corresponding to two thirds of the total leaf number from the bottom were tagged and termed the *n*-th leaf. For transcriptomic studies, we selected the n+1 leaves. Leaves were cut without their petiole and immediately frozen in liquid nitrogen (greenhouse). Grinding was performed using a ZM200 grinder (Retsch, Haan, Germany) with a 0.5-mm sieve. Total RNA was extracted using QIAzol Lysis Reagent following the manufacturer’s instructions (Qiagen, Dusseldorf, Germany). The RNA quality was checked by electrophoresis on an agarose gel and quality and quantity were assessed using the Agilent RNA 6000 nano kit (Agilent, Santa Clara, CA, USA). Sequencing was performed on the Illumina HiSeq 2000 by DNAVision (Charleroi, Belgium) as paired-end libraries (2*100 bp, oriented) using the TruSeq sample preparation kit (Illumina, San Diego, CA, USA) according to manufacturer’s instructions.

The transcriptomic study was performed with the EdgeR package version 3.16.5 (Robinson et al., 2010) on R version 3.3.3 (Another Canoe) (R Core Team, 2017). Genes were considered expressed if there was at least 3 libraries (out of the 142 in the entire experimental design) with at least 2 Counts Per Millions (CPM). CPM were computed with the cpm function from the edgeR package. We found 27,279 genes with at least 2 CPM in 3 libraries. Count normalization was performed as described in the edgeR user guide with the calcNormFactors function using the TMM method (Trimmed Mean normalization). Unless stated otherwise, subsequent computation and analysis were performed on the filtered and normalized values.

To identify Differentially Expressed Genes (DEG) between SF193 and SF193xSF326, the generalized linear model pipeline from edgeR was used as described in the edgeR user manual. Contrasts were created using the makeContrasts function, with the genotypic interaction beeing calculated as: “(SF193xSF326.stress - SF193xSF326.ctrl) - (SF193.stress - SF193.control)”. We corrected p-values using the FDR method and the cutoff was set at 0.05.

### Crop modeling and simulation

#### Model parameterization and evaluation

We used crop modeling and simulation to predict the grain yield in non-observed field environments of genotypes phenotyped in the Heliaphen platform. SUNFLO is a process-based simulation model for sunflower that was developed to simulate grain yield and oil concentration as a function of time, environment (soil and climate), management practice and genetic diversity (Casadebaig et al., 2011; Lecoeur et al., 2011). Predictions with the model are restricted to obtainable yield (Van Ittersum and Rabbinge, 1997): only the main limiting abiotic factors (temperature, light, water and nitrogen) are included in the algorithm.

The model simulates the main soil and plant functions: root growth, soil water and nitrogen content, plant transpiration and nitrogen uptake, leaf expansion and senescence and biomass accumulation. Globally, the SUNFLO crop model has about 50 equations and 64 parameters: 43 plant-related traits, among which eight are genotype-dependent, and 21 environment-related. A report that summarizes the equations and parameters used in the model is available as supplementary information in Picheny et al. (2017). The source code is available on INRA software repository [https://forgemia.inra.fr/record/sunflo.git]. The INRA VLE-RECORD software environment (Bergez et al., 2013; Quesnel et al., 2009) was used as simulation platform.

The model inputs are split in four categories: genotype, climate and soil, management actions and initial conditions.

In the model, a genotype is represented by a combination of eight genotype-dependent parameters whose values are assumed to be constant among environments, thus trying to mimic genetic information (Boote et al., 2003; Jeuffroy et al., 2013). All parameter, except two, were measured directly on field trials (methods in Casadebaig, Mestries, et al., 2016). The two remaining parameters, the responses of leaf expansion and stomatal conductance to water deficit, necessitate controlled-condition experiments. In this study, the measurement protocol for the Heliaphen platform was the same than in greenhouse experiments (Casadebaig et al., 2008). Our goal here was to evaluate if this new phenotyping condition would have an effect on the prediction accuracy of the simulation model.

Four climatic variables are used as daily inputs for simulation: mean air temperature (°C, 2m height), global incident radiation (MJ.m^−2^), potential evapotranspiration (mm, Penman–Monteith) and precipitation (mm). Soil properties are defined by its texture, depth and mineralization, while initial soil conditions are defined by residual nitrogen and initial water content. Needed management practices includes sowing date, planting density, irrigation and nitrogen fertilization.

The SUNFLO simulation model was evaluated both on specific research trials (40 trials, 110 plots) and on agricultural extension trials that were representative of its targeted use (96 trials, 888 plots). Over these two datasets, the model was able to simulate significant genotypes-environment interaction and rank genotypes (Casadebaig et al., 2011; Casadebaig, Mestries, et al., 2016). The prediction error for grain yield was 15.7% when estimated over all data (9% - 30% in individual trials). From these two evaluations, we determined that the model was accurate enough to discriminate between two given genotypes.

#### Impact of parameterization method on model accuracy

Here, we evaluate how the model accuracy is impacted by using the Heliaphen phenotyping platform to estimate the value of two genotype-dependent parameters, the response of leaf expansion and plant transpiration to water deficit. In the parameterization dataset, three genotypes were common between two outdoor experiments and six greenhouse experiments (Casadebaig et al., 2008).

The test dataset was a subset of these three genotypes from a multi-environmental network (described in section plant material) were grain yield was observed in 10 locations. We assessed the model prediction capacity using the bias, the root of mean square error (rmse), and relative rmse, which are different metrics commonly used to evaluate the predicting ability of models. We compared the model prediction capacity on the test dataset, for two sets of simulations: one where genotypes were parameterized with the reference method (greenhouse experiments) and one with parameters measured with the Heliaphen phenotyping platform.

#### Simulation of genotypes in a European trial-network

We created a factorial design from 4 commercial hybrids with contrasted response traits, 42 locations sampled from sunflower cropping area in continental Europe, and 21 years of historical climate data. Genotypes were chosen according to their contrasted responses for leaf expansion and transpiration to water stress, based on data from trials 14HP09 and 15HP04 (Table 1.). We selected a genotype with a rapid response of transpiration and leaf expansion to water stress (*MAS 86OL*), a genotype with a rapid response of transpiration and a slow response of leaf expansion (*LG 5450HO*), a genotype with a slow response of transpiration and rapid response of leaf expansion (*MAS 89M*) and a genotype with slow responses to water stress for transpiration and expansion (*SY EXPLORER*).

The data related to soil properties were obtained by local soil analysis for 13 of 42 trials and completed with the European soil database (ESDAC, esdac.jrc.ec.europa.eu, Panagos et al. (2012); Hiederer (2013)). The climatic dataset was provided by the Agri4cast data portal (Gridded Agro-Meteorological Data in Europe). For each considered location in the study, meteorological data from the nearest grid (25×25 km) point was used, over the 1996-2016 period.

For each genotype × location × year combination (n=3528), six output variables were simulated by the crop model: seed yield at harvest, oil content at harvest, and four time-series of the impact of selected abiotic stress on the simulated photosynthesis rate (water stress, nitrogen deficiency, low temperature stress and high temperature stress).

#### Clustering of environmental time-series

To group environments (location × year combinations) displaying similar abiotic stress patterns, each simulated time-series was summarized by integrating the value of the considered abiotic stress variable over the crop cycle, thus defining a vector of four stress indicators. Water stress indicator was described as the integration of the fraction of transpirable soil water over time (SFTSW). Nitrogen deficiency indicator was defined by the integration of the nitrogen nutrition index (Lemaire and Meynard, 1997) (SNNI). Low and high temperature stress indicators (SLT and SHT) were defined by the integration of photosynthesis response curve to temperature over time. We then used the HCPC function of factomineR (Husson et al., 2010) to make a hierarchical classification of environments based on the simulated stress indicators (3528 environments × 4 indicators).

## Results

### Phenotyping for plant breeding

#### Drought stress management on Heliaphen

The Heliaphen platform aim to precisely control the water deficit level, on different stages of the crop cycle and for various duration. Taking the 16HP07 experiment on the study of the water stress response as an example, we successfully applied different stress levels (figure 3A) during 40 days (from anthesis to harvest) on the commercial genotype NK KONDI. The impact of integrated drought stress indicator (SFTSW) on thousand kernel weight (TKW), seed weight and plant biomass is illustrated in figure 3B, C and D respectively. When modeling the phenotype response to water deficit using the following model 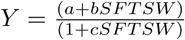, the relative error (rRMSE) was 4.06, 0.20 and 0.07% respectively.

**Figure 3.**
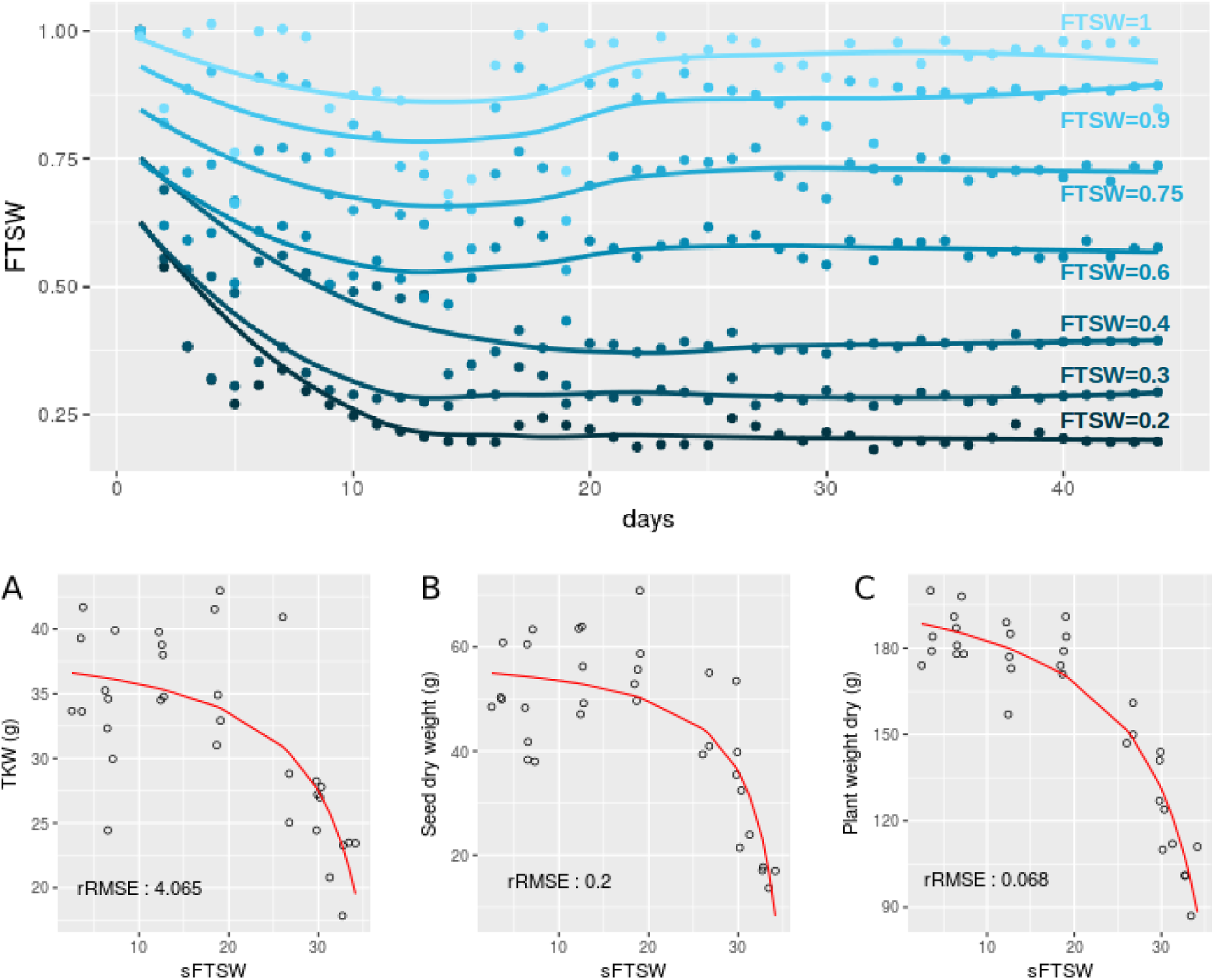
Long-term daily monitoring of water status and impact on yield and biomass traits using the Heliaphen platform. Plants of NK KONDI sunflower genotype were exposed to different water statuses expressed as the Fraction of Transpirable Soil Water (FTSW) after anthesis (day 0) at different thresholds (form dark to light blue corresponding respectively to: 0.2, 0.3, 0.4, 0.6, 0.75, 0.9, 1 of FTSW). Points correspond to the average FTSW on the six plants by stress and by day. Lines correspond to the fit on points by stress with the loess regression method (upper panel). (A) Thousand kernel weight (TKW) response to SFTSW (integration of 1-FTSW) for each plant. (B) Seed weight (per plant) response to SFTSW. (C) Plant weight response to SFTSW. rRMSE: relative root mean square error of the polynomial ratio model.

#### Comparison between phenotypes observed on Heliaphen platform and in field experimentation

We also evaluated how traits measured on the platform environment were related to field conditions. For this purpose we compared the weight of seeds per plant obtained on trials conducted on the platform (14HP10 trial) and in field (14RV01 trial), which shared the same genetic material, growing period and environmental conditions. The seed weight measurements between the two tests were significantly correlated (R = 0.23, p-value < 0.001, figure 4B). This panel was also evaluated in eight other trials in a European network spanning over three years (trials 13EX01 to 15EX07). The average correlation between the field trials is 0.22 as shown in figure 4B, which is similar to the correlation level found between platform and field trial grown simultaneously (14HP10 and 14RV01). The Heliaphen trial was also significantly correlated to two other field trials (14EX04 R=0.17 and 15EX07 R=0.13).

**Figure 4.**
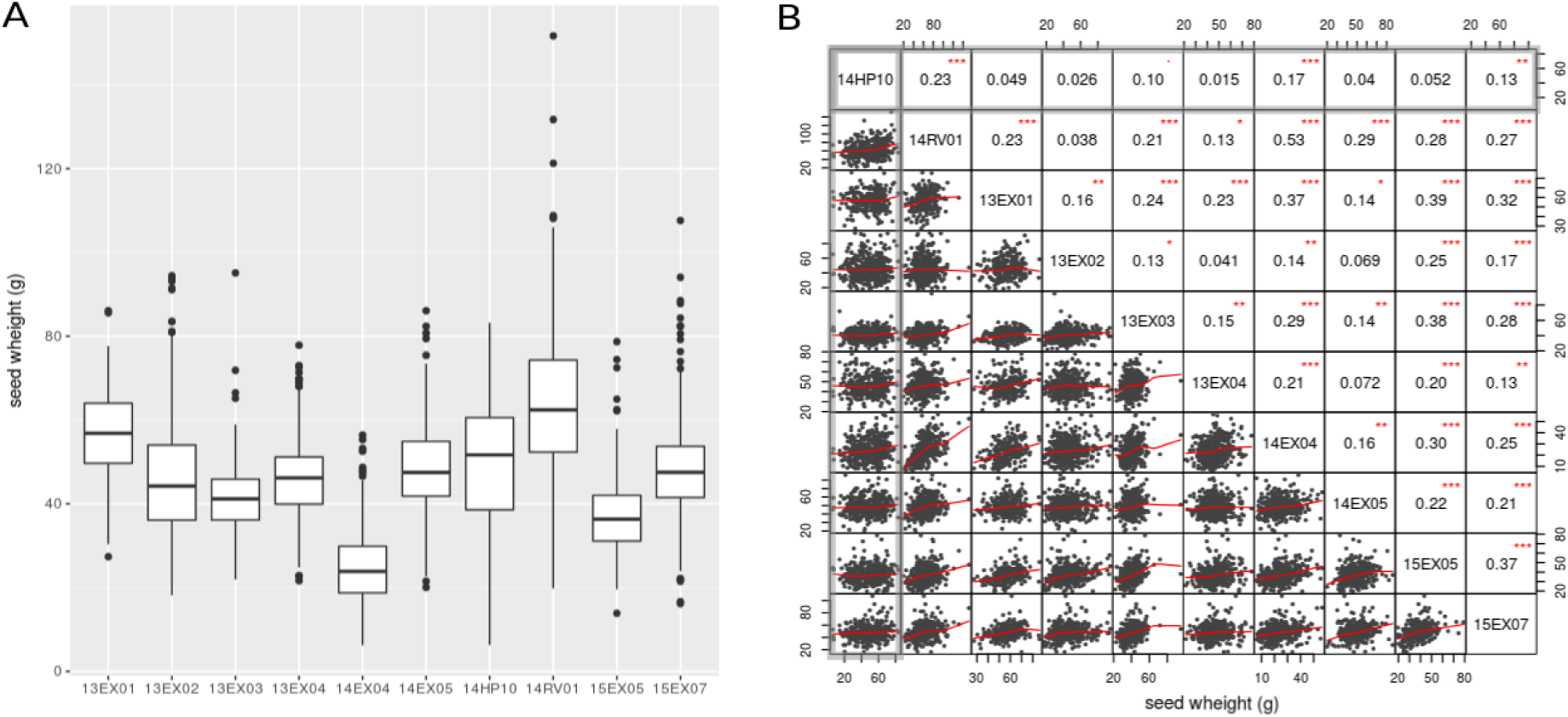
Comparison between seed mass per plant measured on the Heliaphen phenotyping platform and in field conditions. (A) Boxplot of the seed dry weight observed on the SUNRISE panel. (B) Correlation matrix between plant seed dry weight observed on the SUNRISE hybrid panel (n= 426) on the platform (14HP10 highlighted in gray box) and ten field trials (14RV01 at 15EX07). Correlation coefficients are on the top right part of the figure with significance test indicated with stars (* corresponds to 0.05 threshold, ** to 0.01 and *** to 0.001)

**Figure 5.**
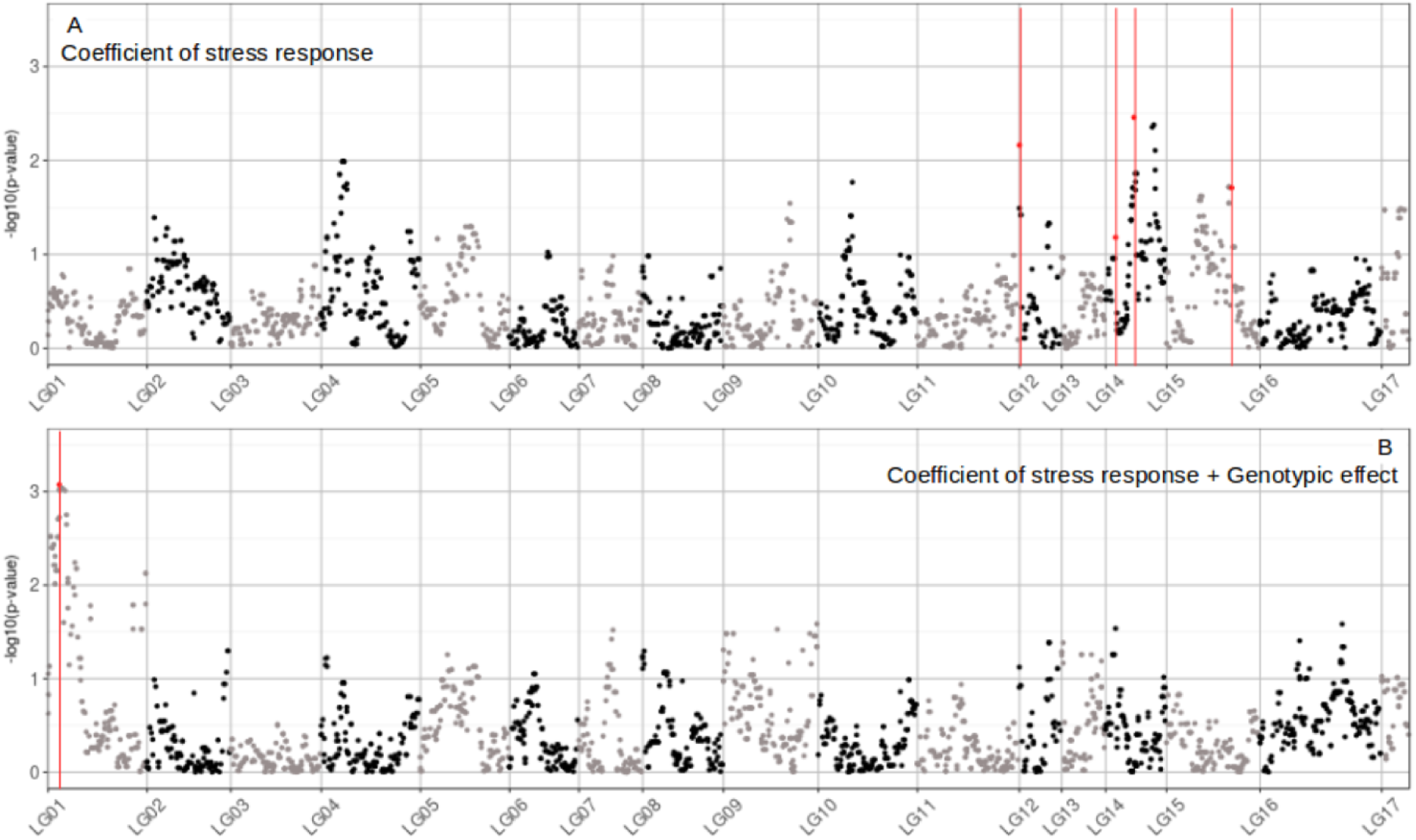
Manhattan plot of the GWAS performed two traits related to water stress tolerance. (A) GWAS on the genetic coefficient of stress response. (B) GWAS on the sum of the coefficient of stress response and the genotypic effect (total genetic effect). The markers found associated by MLMM method are highlighted in red. The p-value on the graph corresponds to that obtained with a model of classical GWAS, performed during the first step of the MLMM method.

Figure 4A shows that although pedoclimate has a strong impact on seed production of our panel, and that we observe significant differences between the variances and the means on the different trials, the seed weight distribution measured on the platform is in the same range than the field trials.

#### Association genetics on phenotypes measured on the Heliaphen platform

We performed a genome wide association study on the response of seed weight to water deficit (constant water deficit on reproductive phase, FTSW = 0.4), using a RILs population (13HP05 trial). The thousand kernel weight response was modeled using a linear model of water stress indicators (cf. the genetic study section of the Material and Methods). In order to identify markers associated with the response to water stress, a association test (MLMM) was performed on the genetic coefficient of stress response (*α_i_*, interaction effect) and on the sum between the stress response coefficient and the genetic effect (*γ_i_* + *α_i_*, total genetic effect). In this approach, we selected the set of markers that together explain the genetic variance of the studied characters. We obtained four markers associated to the coefficient of response to the water stress and one to the sum of the stress response coefficient and genetic effect (Table 2). All markers were positioned on the sunflower genetic map and the Manhattan plot on figure 4 allows to visualize the p-value calculated for the first MLMM step corresponding to classical model of GWAS (Yu et al., 2006).

**Table 2.** Markers associated to response to drought stress of thousand kernel weight and candidate genes showing different response to drought in the RIL population parents from transcriptomics analysis). Positions, names and distances are based on the HanXRQv1 genome.

| Trait | Marker | LG | Position (bp) | Effect | Gene | Distance (bp) |
| --- | --- | --- | --- | --- | --- | --- |
| $\alpha_i$ | AX-105774006 | 12 | 164538019 | 0.08 | HanXRQChr12g0385481 | 1258962 |
| $\alpha_i$ | AX-105362253 | 14 | 123649962 | 0.12 | HanXRQChr14g0445121 | 1443617 |
| $\alpha_i$ | AX-105376351 | 14 | 167871915 | -0.11 | HanXRQChr14g0459681 | 20774 |
| $\alpha_i$ | AX-105568999 | 15 | 92795704 | 0.08 | HanXRQChr15g0486311 | 2980461 |
| $\alpha_i + \gamma_i$ | AX-105798220 | 1 | 61598886 | 2.74 | HanXRQChr01g0009921 | 4628392 |

#### Transcriptomics analysis

In order to focus on the best candidate genes carrying the causal polymorphisms, we studied gene expression in both homozygote SF193 and heterozygous SF193/SF326 backgrounds, identical to the genetic design used for association. Based on the hypothesis that the causal polymorphism acts on gene expression, we looked for the gene showing the most significant interaction between drought stress and genotype around the associated markers (± 5Mb) in the transcriptomic measurements performed on the 13HP02 trial (figure 6).

**Figure 6.**
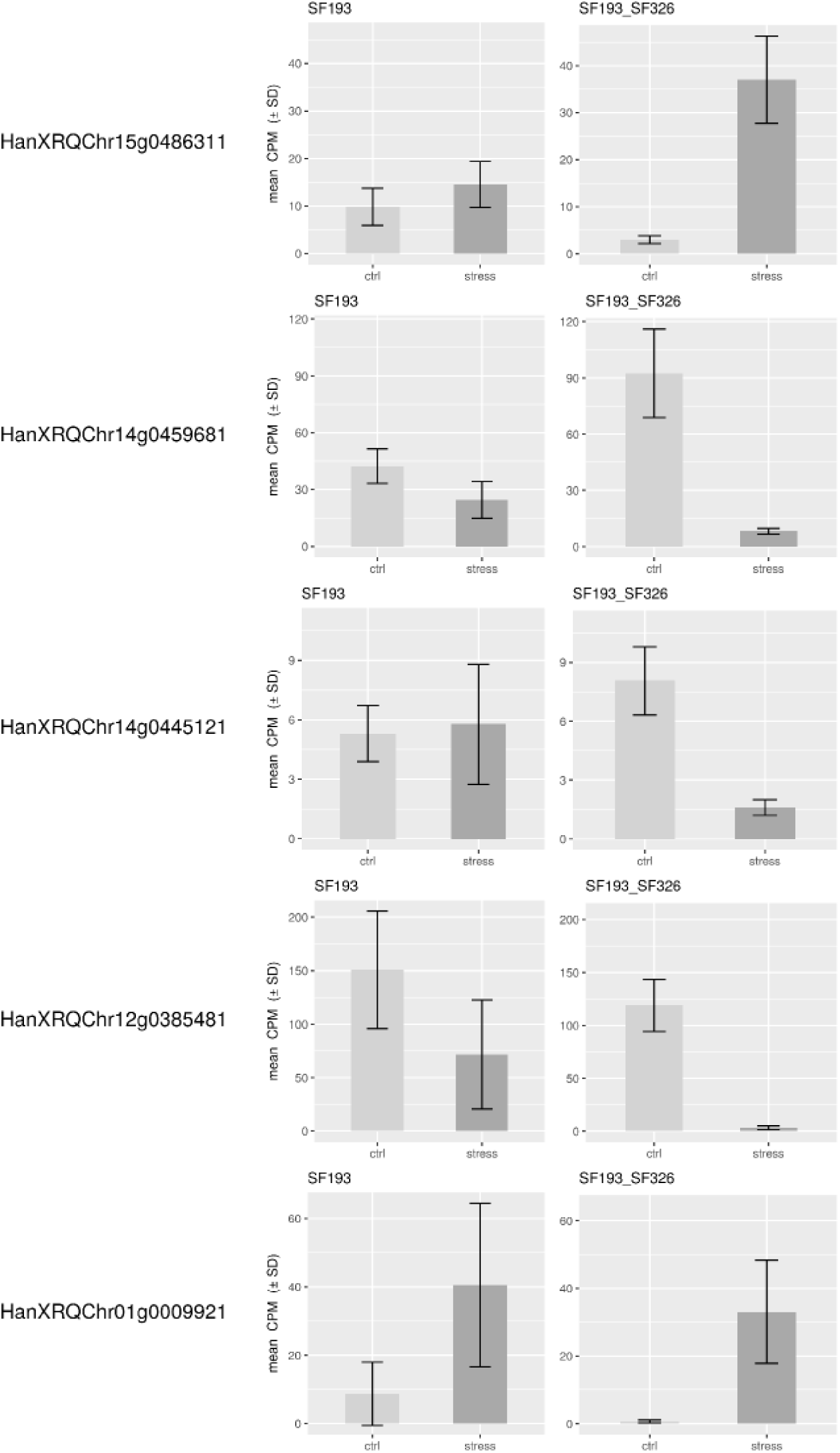
Expression levels of genes related to associated SNPs. Expression levels of genes with the most differentially expression for the genotype-drought interaction at ± 5Mb to the associated SNPs to drought stress response in the back-crossed RIL population (SF193x(SF193xSF326)). Expression levels (Count Per Million, CPM) were evaluated in water stress condition (stress, dark grey) or normal watering condition (ctrl, light grey), in SF193 and SF193xSF326.

On chromosome 12, we identified an associated marker to the drought stress coefficient. The effect of the alternative allele of SF326 on the coefficient of stress response was 0.084, providing drought tolerance. Following this approach, we were able to identify the gene HanXRQChr12g0385481. The marker is situated 1.2Mb downstream of the gene. This gene is homologous (54% of protein identity) to the Arabidopsis protein UDP-GLYCOSYLTRANSFERASE 90A1 (UGT90A1) and show homology with proteins belonging to the family of UDP-glycosyltransferase including among others UGT79E11. The family of UDP-glycosyltransferase is a wide protein family and some of them are involved in abiotic stress (including drought) responses in Arabidopsis (Pan Li, Y.-j. Li, et al., 2016; Pan Li, Y.-J. Li, et al., 2016; Li et al., 2017).

On chromosome 14, we identified two associated markers to the drought stress coefficient. The effects of the alternative allele SF326 were 0.118 and −0.106 providing tolerance and sensitivity respectively. In the same way than the marker found associated on the chromosome 12, we studied gene expression. We identified the gene HanXRQChr14g0445121, its marker is situated 1.4Mb downstream of the gene. This gene is homologous (47.71% of protein identity) to the Arabidopsis protein RING-H2 FINGER PROTEIN ATL54, a gene involved in the secondary cell wall formation (Noda et al., 2013). The second gene identified on the chromosome 14 is HanXRQChr14g0459681, situated 20kb to its marker (downstream of the gene). This gene is homologous (38.45% of protein identity) to the Arabidopsis protein SMAX1-LIKE 7 (SMXL7) involved in the regulation of the shoot development (Liang et al., 2016).

On chromosome 15, one marker has been identified associated to the drought stress coefficient. The effect of the alternative allele SF326 was 0.085 providing tolerance. We identified HanXRQChr15g0486311, this gene is situated 2.9Mb to its marker (downstream of the gene). This gene is homologous with two Arabidopsis transcription factors TCP19 (52.81 % of protein identity) and TCP9(46.80% of protein identity). These transcription factors are involved in the control of leaf senescence and the regulation of the Jasmonic Acid (Danisman et al., 2013, 2012) that were largely documented to be linked to drought stress response in different species including sunflower (Andrade et al., 2017; Marchand, 2014).

On chromosome 1, one marker has been associated to total genetic effect (sum of the drought stress genetic coefficient and genetic effect). The effect of the alternative allele SF326 was 2.737, providing tolerance. We identified the gene HanXRQChr01g0009921, situated at 4.6Mb to its marker (downstream of the gene). This gene is homologous (66.47% of protein identity) to the Arabidopsis protein monofunctional riboflavin biosynthesis protein RIBA3, a protein involved in the synthesis of riboflavin (Hiltunen et al., 2012) that are suggested to alleviate ROS production during osmotic stress (Deng et al., 2013).

The Heliaphen platform successfully allowed us to identify genomic regions associated to a water stress response for yield traits. Using this information, we could confront it to other data and identify candidate genes putatively involved in the response to water stress. Further studies are needed to prove their involvement in the response of yield to water stress, but the platform was instrumental to generate these promising results.

### Phenotyping for crop modeling

#### Impact of parameterization method on the crop model accuracy

The SUNFLO crop model (Casadebaig et al., 2011) use phenotypic traits as genotype-dependent input parameters. Among them, two parameters describe the response of leaf expansion and transpiration to water stress. The measurement method requires dynamic leaf area measurements and assessment of water deficit at the plant level. Here, we evaluate how the measurement of these parameters with the Heliaphen platform impacts the model accuracy as compared to the reference method based on greenhouse experiments (Casadebaig et al., 2008).

We first compared the standard error of the parameters estimations performed with measurements in either Heliaphen or greenhouse conditions. Parameters were measured for commercial hybrids genotypes and are presented in figure 7. Lower parameter values indicate the genotypes start to regulate either leaf expansion or transpiration at lower FTSW values, indicating that the genotype maintain carbon assimilation at lower water deficit. We observe experimentations on the Heliaphen platform resulted in similar parameter estimation error for transpiration response (~ 1.1 sd for both conditions) and halved error for leaf expansion response (0.3 sd Vs 0.7 sd in greenhouse). Additionaly, the estimation error was not dependent of the parameter value, as it was the case in greenhouse. Because these traits are used as input values in the simulation model and have an impact on yield prediction, we also evaluated how the model prediction accuracy is impacted when using different parameterization conditions. We used the relative root mean square error (RRMSE), the ratio of the root mean square error on the average yield. The RRMSE on the yield prediction were equivalent (13 %) between the simulations made from the response traits obtained from the Heliaphen platform trials and those from the greenhouse trials.

**Figure 7.**
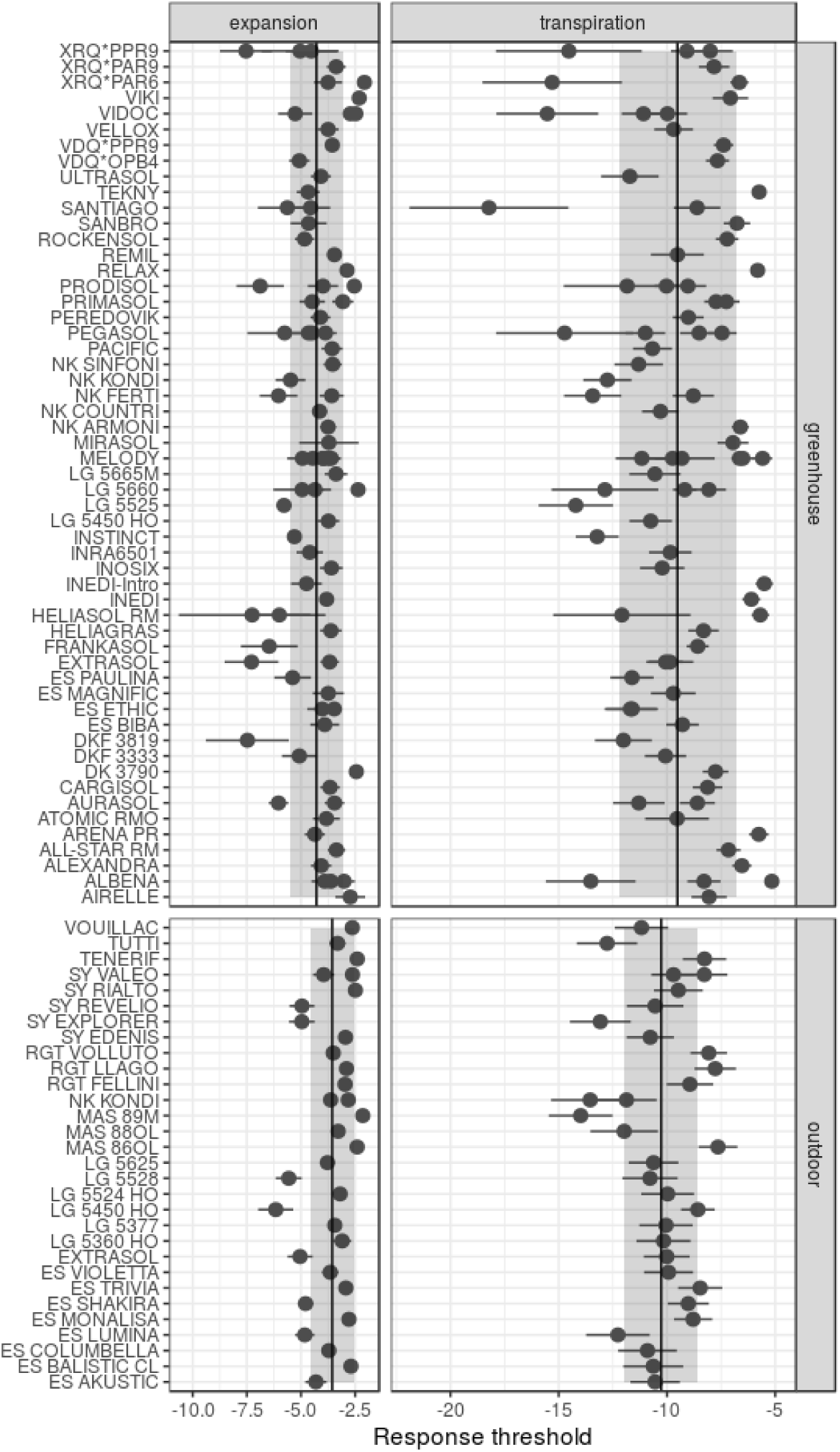
Drought response parameters of the SUNFLO crop model measured in greenhouse environment and on Heliaphen platform, for different commercial hybrids. Points correspond to the value of parameter for each genotype. Ranges indicated with lines correspond to the standard error of estimated parameter, ranges indicated with a shaded area correspond to the standard deviation for the population phenotyped in greenhouse or Heliaphen condition.

We can conclude that the prediction accuracy is conserved with the measurements on the Heliaphen platform and that this platform facilitates the measurement of these two input parameters while achieving a greater precision.

#### Integrating traits measured with the Heliaphen platform to simulate genotype-environment interactions

The Heliaphen platform allows to measure the genotype-dependent traits of varieties. This information can be fed into a simulation model to estimate their productivity in different types of environments. To illustrate this approach, we selected four genotypes with contrasting water stress response traits (Table 3), used a computational experiment to compare them in a trial network representative of sunflower growing regions in Europe. The selected genotypes were simulated with the SUNFLO crop model on 42 locations (figure 8C) over 21 climatic years from 1996 to 2016.

**Figure 8.**
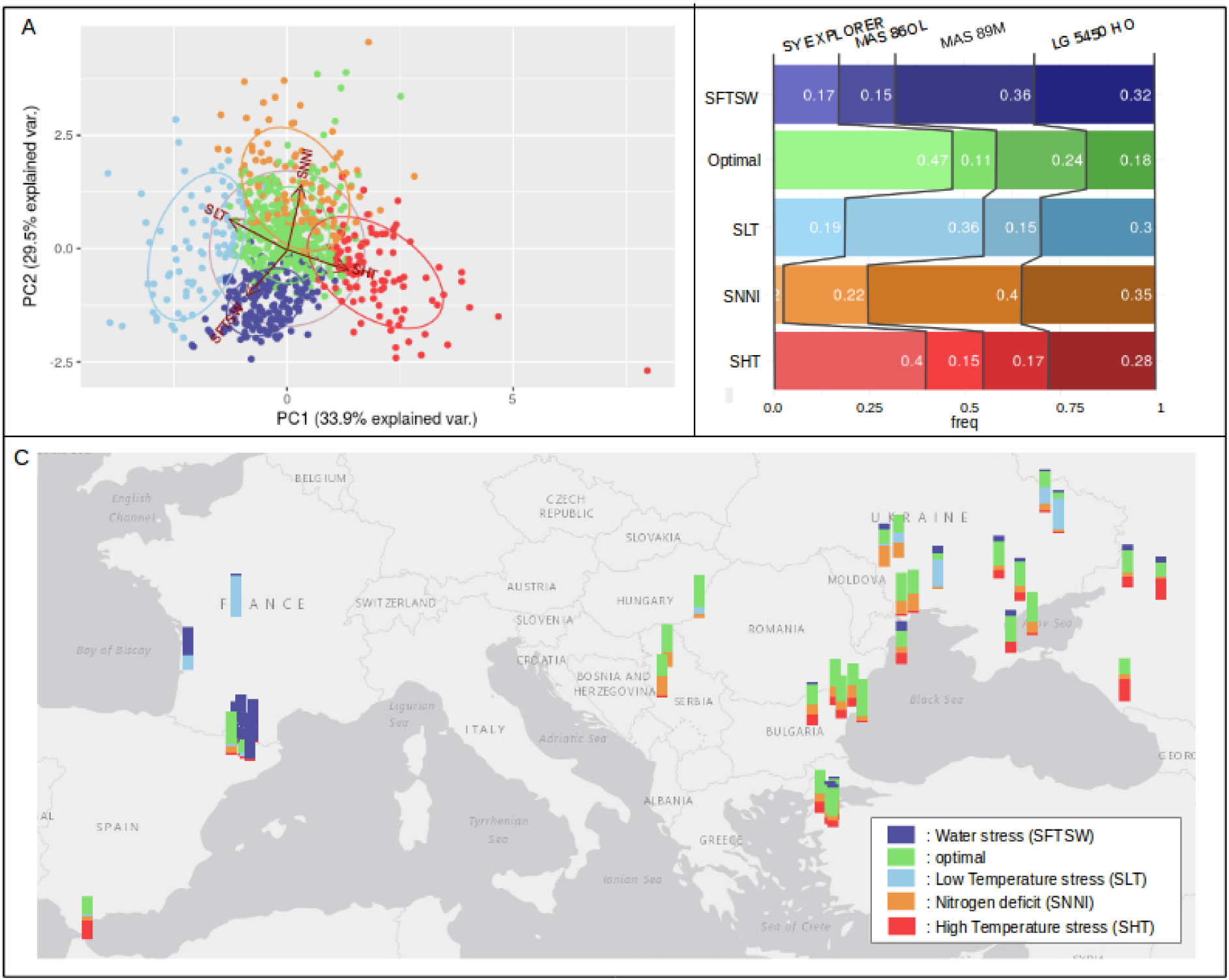
Use of simulation modeling to characterize cropping environments. Description of stress frequencies in 42 sunflower production areas at the European scale. (A) Clustering of environments (year-location) on the principal components of average simulated stresses for four sunflower genotypes. The PCA was performed using High Temperature stress (SHT), Low Temperature stress (SLT), Water stress (SFTSW) and Nitrogen deficit (SNNI). Variables are represented with red arrows. Clustering is on five groups: four associated to a specific stress and named accordingly and the one without obvious stress was named Optimal. (B) Standardized yield of each genotype have been ranked on simulation group by environment, the genotype frequency at the first rank in each cluster is reported inside the bars. (C) Map of Europe with a barplots at each location. Colors on barplots indicate the occurrence of the location in the different environment clusters.

**Table 3.** Response parameters of genotypes used for simulation, (+) means fast response to water stress compare to other genotype (fig 7) and (-) means slow response.

| Genotype | Transpiration rate | Leaf expansion rate |
| --- | --- | --- |
| LG 5450H0 | - 9,66 (+) | -4,94 (-) |
| MAS 86OL | -7,64 (+) | -2,40 (+) |
| MAS89M | -13,98 (-) | -2,15 (+) |
| SY EXPLORER | -13,08 (-) | -4,96 (-) |

We defined an environment as the combination of a location and a year, and we generated simulations for each genotype on 882 environments (42 locations × 21 climatic years). On these 882 environments, we calculated the average of the abiotic stress indicators for the four genotypes. Figure 8A is a PCA constructed from average stress indicators, where each point is an environment. Five environment clusters could be defined and each group (or environment-type, Chenu et al., 2013) was labeled following the projection of stress indicators in the PCA plane. The relative frequencies of each environment-type could then be mapped to the corresponding geographical locations (figure 8C) to illustrate the most frequent abiotic stress factors in the different sunflower-growing regions across Europe. For example, the high frequency of drought stress in France, could be explained by the impact of the shallower soil depth in these locations.

We also aimed to identify environment-types were specific genotypes performed better, relatively to the other genotypes tested. For that, we adjusted the simulated yields by the average yield of each genotype and then ranked the genotypes on this index in each environment. Figure 8B shows, for each environment-type, the frequency of the best ranking for each genotypes. Thus, we can characterize the abiotic stress tolerance profiles for the genotypes, or said in another way, what kind of environment maximizes the yield of each genotype compared to the other. Based on this method, we could show that LG5450HO displayed an equilibrated tolerance profile (good frequency for each environment-type), SY EXPLORER was particularly adapted to heat-stressed environments, and MAS89M to nitrogen-stress and water-stressed ones. In this approach, the Heliaphen platform was a key step to characterize genotypes to better select them and identify the areas where they could potentially perform best.

## Discussion

In order to characterize genotypes for drought stress response, we developed the Heliaphen high-throughput phenotyping platform. This facility allows different water stress scenarios to be applied to plants, to evaluate the effect of these stresses on yield and its components. Results obtained on the platform confirmed its reliability to implement water stress on large panels of plants. Additionally, specific plant traits can be measured and used as input parameters of crop simulation models.

The Heliaphen platform is a unique phenotyping facility that stands out against other methods of automating phenotyping. Most phenotyping platforms rely in strongly controlled environmental conditions and are operated inside greenhouse or growth chambers (Cabrera-Bosquet et al., 2016; Granier et al., 2005; Honsdorf et al., 2014; Pereyra-Irujo et al., 2012) and mainly target the vegetative part of the plant cycle, mainly due to the size and space that a mature crop plant takes. These facilities make it possible to control the biotic and abiotic environment more finely but consequently move away from the field conditions differing by the light spectrum and atmospheric conditions. On the contrary, phenotyping tools operating in field conditions (Unmanned Aerial Vehicles) (Hu et al., 2018; Sankaran et al., 2015) or robots (Deswarte et al., 2015; Underwood et al., 2017) measure plants in actual agricultural conditions with little control over climate scenarios.

The automation of phenotyping tools is continuously under development on the platform. Currently on-going developments for sunflower involve using different sensors type to access new phenotypes: ultrasonic sensors (plant height), laser scanner (stem diameter), and a light curtain sensor (leaf area profile, Rapidoscan RS-C-025-768-ECT). For exemple, the temporal analysis of organs (stem, petiol, leaf blades) growth was greatly improved by developing an image analysis algorithm. In this method, a 3D point cloud is computed using Structure from Motion techniques based on RGB images acquired around the plant. The 3D cloud is then segmented into organs whose area, size and volume can be estimated (Gélard et al., 2017). In addition to sunflower, we use the platform to monitor the growth and development of other species such as soybeans, corn, and tomatoes. It would also be possible to conduct trials combining several stresses, to study for example how nutrient stress interact with water deficit.

For the Heliaphen facility, we first wanted to validate the possibility to design specific drought scenarios and compared yield-related traits assessed on the platform to field conditions. We successfully applied different stress intensities over periods of various length, such as the complete grain filling period (about 40 days). We confirmed our ability to relate the water deficit level with its impact on yield components (such as seed number and weight). In addition, the comparison of seed production between field trials and a platform trial showed a correlation (quite poor, but highly significant thanks to the power of the statistical design) for seed weight comparable with those observed between two field trials. This highlights on one hand the importance of genotype-environment interactions and the difficulty to predict them, and on the other hand the major interest of the facility to produce phenotypes relevant in agronomic conditions. It is therefore possible to study and identify genes controlling drought tolerance for yield.

### Identify genes controlling drought tolerance for yield

From a RIL population subjected to different scenarios of water stress, we conducted a genetic study and identified the regions associated with the response of seed weight to water stress. Five markers have been identified. The analysis of gene expression in stressed and unstressed conditions was conducted in both genetic backgrounds. Following the hypothesis that the causal polymorphisms act on gene expression, we looked for the most differentially expressed genes in the region surrounding the identified markers and studied them as potential candidates. The results are very encouraging, among the identified genes, several stand out. There is a protein belonging to the UDP-glycosyl transferase family, a large family of proteins, some of which are involved in the responses to abiotic stress (including drought) in Arabidopsis (Pan Li, Y.-j. Li, et al., 2016; Pan Li, Y.-J. Li, et al., 2016; Li et al., 2017). We also detected a putative TCP transcription factor homologous to AtTCP19 involved in the control of leaf senescence and regulation of jasmonic acid metabolism (Danisman et al., 2013, 2012) that were largely documented to be linked to drought stress response in different species including sunflower (Andrade et al., 2017; Marchand, 2014).

Another candidate gene is encoding a protein homologous to riboflavin RIBA3, a protein involved in the synthesis of riboflavin (Hiltunen et al., 2012) that is suggested to relieve the production of ROS during osmotic stress (Deng et al., 2013). However, the distance between the identified genes and the position of the markers is sometimes very large (the marker detected on chromosome 1 is more than 4Mb of the gene). The association markers represent genomic regions of variable extension between two recombinations. So it is not surprising to have a large distance between the markers and the differentially expressed genes even if we must pay attention to the trust we place in the choice of genes candidates. Choosing genes differentially expressed in dry conditions positioned near the markers associated to drought response, constitutes an efficient method to identify a small number of relevant genes of interest.

The previously used method could be applied to identify markers or genes associated with other water stress scenarios. We can hope to identify the genomic regions associated with the water stress response applied on other periods and organs, such as the vegetative period or later on developing seeds, and combine this with transcriptomics to identify genes associated with stress response and development under stress conditions. Of course, the platform also allows functional studies such as the transcriptomic analysis (Badouin et al., 2017) that allowed us to identify differentially expressed genes under water stress conditions and other omics combined with manual classical physiological descriptions (photosynthesis, transpiration, osmotic adjustment, etc…).

### Using simulation to predict genotype-environment interactions

The other major goal of the platform was to integrate ecophysiology and crop modeling. Thanks to the simultaneous monitoring of water deficit level, leaf area and water loss dynamics, we showed that physiological traits measured on the platform were more precise than in greenhouse and successfully used as parameters in a crop simulation model. The accuracy of the SUNFLO model was similar with the reference greenhouse-based measurements or those obtained on the platform. The platform setup saves both time and pain compared to previous set up with manual greenhouse measurements and therefore allows us to evaluate the stress response traits of a large number of genotypes each year, keeping pace with the newly bred varieties. We also wanted to expand the scope of our study by designing a simulation experiment were the phenotyped genotypes are used to reveal abiotic stress occurring at the plant level, in a large range of environments. Similar cropping condition are then grouped together to identify broad environment-types were stress dynamics are more predictable (also known as envirotyping, Xu, 2016). Using this method, we identified the major stresses occurring on the various sampled locations.

The accuracy of this method depends on the uncertainty of data used to describe the locations (climate, soil properties, crop management systems). Even if soil and climate databases exists, we found that the method was particularly sensitive to the estimation of soil available water capacity.

With finer environmental descriptions available in the future, we hope to increase the resolution and span of the simulations. This would be a key step to extend classical variety evaluation, and work on recommendation zones based on environment-types identified with simulation studies. Although the variety recommendation method performed in this study was illustrated for abiotic stress tolerance and cannot yet constitute an operational choosing method for growers (as genotypes can greatly differ for their potential yield), this is an important step to link plant phenotyping platforms and model-based recommendation of varieties as performed previously (Casadebaig, Mestries, et al., 2016). In addition, model-based environmental characterization was also showed powerful for genetic study of yield response to combined abiotic stresses (Mangin et al., 2017). This original approach showed that simulated variables related to abiotic stress were closer (more related to yield) to the plant stress and therefore gene action rather variables derived from pedo-climatic data (Lasky et al., 2015; Rasheed et al., 2017).

A challenge for plant breeders is to predict the performance of novel genetic material in different pedo-climatic environments and with different managements. To achieve this, predictive approaches bridging quantitative genetics and crop modeling (Bustos-Korts et al., 2016) are under development to scale traits from the molecular to the crop level. One option is to process in two steps: first to use genomic prediction to estimate genotype-dependent parameters (traits assumed to be more heritable) of a crop simulation model and then use the model to predict performance-related traits (assumed to handle genotype-environment interactions) as a function of environmental data (e.g. Chenu et al., 2009 for maize; Quilot-Turion et al., 2016, for peach tree). The first step, i.e. the genetic analysis of the input parameters of the model either to estimate allelic effects of QTL or to train a genomic selection training population largely rely on phenotyping capacity (Cooper et al., 2014). Such analysis can be done using the Heliaphen platform and illustrates its high potential to accelerate the understanding of genetic control of drought stress plasticity, characterize complex response traits of genotypes. It also enables crop model based approaches to characterize and cluster cropping environments, leading to a better fitting of genotypes in their optimal growing conditions.

## References

Adiredjo, A.L., Navaud, O., Grieu, P., Lamaze, T., 2014. Hydraulic conductivity and contribution of aquaporins to water uptake in roots of four sunflower genotypes. Botanical studies 55, 75. https://doi.org/10.1186/s40529-014-0075-1

Adiredjo, A.L., Navaud, O., Muños, S., Langlade, N.B., Lamaze, T., Grieu, P., 2014. Genetic control of water use efficiency and leaf carbon isotope discrimination in sunflower (helianthus annuus l.) subjected to two drought scenarios. PloS one 9, e101218. https://doi.org/10.1371/journal.pone.0101218

Andrade, A., Escalante, M., Vigliocco, A., Carmen Tordable, M. del, Alemano, S., 2017. Involvement of jasmonates in responses of sunflower (helianthus annuus) seedlings to moderate water stress. Plant Growth Regulation 83, 501–511. https://doi.org/10.1007/s10725-017-0317-9

Andrianasolo, F.N., Casadebaig, P., Langlade, N., Debaeke, P., Maury, P., 2016. Effects of plant growth stage and leaf ageing on transpiration and photosynthesis response to water stress in sunflower. Functional Plant Biology. https://doi.org/10.1071/FP15235

Andrianasolo, F.N., Champolivier, L., Debaeke, P., Maury, P., 2016. Source and sink indicators for determining nitrogen, plant density and genotype effects on oil and protein contents in sunflower achenes. Field Crops Research 192, 33–41. https://doi.org/10.1016/j.fcr.2016.04.010

Badouin, H., Gouzy, J., Grassa, C.J., Murat, F., Staton, S.E., Cottret, L., Lelandais-Brière, C., Owens, G.L., Carrère, S., Mayjonade, B., Legrand, L., Gill, N., Kane, N.C., Bowers, J.E., Hubner, S., Bellec, A., Bérard, A., Bergès, H., Blanchet, N., Boniface, M.-C., Brunel, D., Catrice, O., Chaidir, N., Claudel, C., Donnadieu, C., Faraut, T., Fievet, G., Helmstetter, N., King, M., Knapp, S.J., Lai, Z., Paslier, M.-C.L., Lippi, Y., Lorenzon, L., Mandel, J.R., Marage, G., Marchand, G., Marquand, E., Bret-Mestries, E., Morien, E., Nambeesan, S., Nguyen, T., Pegot-Espagnet, P., Pouilly, N., Raftis, F., Sallet, E., Schiex, T., Thomas, J., Vandecasteele, C., Varès, D., Vear, F., Vautrin, S., Crespi, M., Mangin, B., Burke, J.M., Salse, J., Muños, S., Vincourt, P., Rieseberg, L.H., Langlade, N.B., 2017. The sunflower genome provides insights into oil metabolism, flowering and asterid evolution. Nature 546, 148. https://doi.org/10.1038/nature22380

Bergez, J., Chabrier, P., Gary, C., Jeuffroy, M., Makowski, D., Quesnel, G., Ramat, E., Raynal, H., Rousse, N., Wallach, D., Debaeke, P., Durand, P., Duru, M., Dury, J., Faverdin, P., Gascuel-Odoux, C., Garcia, F., 2013. An open platform to build, evaluate and simulate integrated models of farming and agro-ecosystems. Environmental Modelling & Software 39, 39–49. https://doi.org/10.1016/j.envsoft.2012.03.011

Blanchet, R., Marty, J.-R., Merrien, A., Puech, J., 1981. Main factors limiting sunflower yield in dry areas, in: Production and Utilization of Protein in Oilseed Crops. Springer, pp. 205–226. https://doi.org/10.1007/978-94-009-8334-2_18

Boote, K.J., Jones, J.W., Batchelor, W.D., Nafziger, E.D., Myers, O., 2003. Genetic Coefficients in the CROPGRO-Soybean Model: Links to Field Performance and Genomics. Agronomy Journal 95, 32–51.

Bustos-Korts, D., Malosetti, M., Chapman, S., Eeuwijk, F. van, 2016. Modelling of genotype by environment interaction and prediction of complex traits across multiple environments as a synthesis of crop growth modelling, genetics and statistics, in: Crop Systems Biology. Springer, pp. 55–82. https://doi.org/10.1007/978-3-319-20562-5_3

Butler, D., Cullis, B., Gilmour, A., Gogel, B., 2009. ASReml-r reference manual. Queensland Department of Primary Industries, Queensland, Australia.

Cabrera-Bosquet, L., Crossa, J., Zitzewitz, J. von, Serret, M.D., Araus, J.L., 2012. High-throughput phenotyping and genomic selection: The frontiers of crop breeding ConvergeF. Journal of Integrative Plant Biology 54, 312–320. https://doi.org/10.1111/j.1744-7909.2012.01116.x

Cabrera-Bosquet, L., Fournier, C., Brichet, N., Welcker, C., Suard, B., Tardieu, F., 2016. High-throughput estimation of incident light, light interception and radiation-use efficiency of thousands of plants in a phenotyping platform. New Phytologist 212, 269–281. https://doi.org/10.1111/nph.14027

Casadebaig, P., Debaeke, P., Lecoeur, J., 2008. Thresholds for leaf expansion and transpiration response to soil water deficit in a range of sunflower genotypes. European Journal of Agronomy 28, 646–654. https://doi.org/10.1016/j.eja.2008.02.001

Casadebaig, P., Guilioni, L., Lecoeur, J., Christophe, A., Champolivier, L., Debaeke, P., 2011. SUNFLO, a model to simulate genotype-specific performance of the sunflower crop in contrasting environments. Agricultural and Forest Meteorology 151, 163–178. https://doi.org/10.1016/j.agrformet.2010.09.012

Casadebaig, P., Mestries, E., Debaeke, P., 2016. A model-based approach to assist variety assessment in sunflower crop. European Journal of Agronomy 81, 92–105. https://doi.org/10.1016/j.eja.2016.09.001

Casadebaig, P., Zheng, B., Chapman, S., Huth, N., Faivre, R., 2016. Assessment of the potential impacts of wheat plant traits across environments by combining crop modeling and global sensitivity analysis. PLoS ONE 11, e0146385. https://doi.org/10.1371/journal.pone.0146385

Chapman, S., Cooper, M., Hammer, G., 2002. Using crop simulation to generate genotype by environment interaction effects for sorghum in water-limited environments. Australian Journal of Agricultural Research 53, 379–389.

Chapman, S., Cooper, M., Podlich, D., Hammer, G., 2003. Evaluating Plant Breeding Strategies by Simulating Gene Action and Dryland Environment Effects. Agronomy Journal 95, 99–113. https://doi.org/10.2134/agronj2003.0099

Chaves, M.M., Maroco, J.P., Pereira, J.S., 2003. Understanding plant responses to drought—from genes to the whole plant. Functional plant biology 30, 239–264. https://doi.org/10.1071/fp02076

Chenu, K., Chapman, S., Tardieu, F., McLean, G., Welcker, C., Hammer, G., 2009. Simulating the yield impacts of organ-level quantitative trait loci associated with drought response in maize: A “gene-to-phenotype” modeling approach. Genetics 183, 1507. https://doi.org/10.1534/genetics.109.105429

Chenu, K., Deihimfard, R., Chapman, S.C., 2013. Large-scale characterization of drought pattern: A continent-wide modelling approach applied to the Australian wheatbelt–spatial and temporal trends. New Phytologist 198, 801–820. https://doi.org/10.1111/nph.12192

Cobb, J.N., DeClerck, G., Greenberg, A., Clark, R., McCouch, S., 2013. Next-generation phenotyping: Requirements and strategies for enhancing our understanding of genotype–phenotype relationships and its relevance to crop improvement. Theoretical and Applied Genetics 1–21.

Cooper, M., Messina, C.D., Podlich, D., Totir, L.R., Baumgarten, A., Hausmann, N.J., Wright, D., Graham, G., 2014. Predicting the future of plant breeding: Complementing empirical evaluation with genetic prediction. Crop and Pasture Science 65, 311–336. https://doi.org/10.1071/cp14007

Danisman, S., Dijk, A.D.J. van, Bimbo, A., Wal, F. van der, Hennig, L., Folter, S. de, Angenent, G.C., Immink, R.G.H., 2013. Analysis of functional redundancies within the arabidopsis TCP transcription factor family. Journal of Experimental Botany 64, 5673–5685. https://doi.org/10.1093/jxb/ert337

Danisman, S., Wal, F. van der, Dhondt, S., Waites, R., Folter, S. de, Bimbo, A., Dijk, A.D. van, Muino, J.M., Cutri, L., Dornelas, M.C., Angenent, G.C., Immink, R.G.H., 2012. Arabidopsis class i and class II TCP transcription factors regulate jasmonic acid metabolism and leaf development antagonistically. PLANT PHYSIOLOGY 159, 1511–1523. https://doi.org/10.1104/pp.112.200303

Debaeke, P., Casadebaig, P., Flenet, F., Langlade, N., 2017. Sunflower and climate change in europe: Crop vulnerability, adaptation, and mitigation potential. Oilseeds and fats, Crops and Lipids. https://doi.org/10.1051/ocl/2016052

Deng, B., Jin, X., Yang, Y., Lin, Z., Zhang, Y., 2013. The regulatory role of riboflavin in the drought tolerance of tobacco plants depends on ROS production. Plant Growth Regulation 72, 269–277. https://doi.org/10.1007/s10725-013-9858-8

Deswarte, J.-C., Beauchene, K., Arjaure, G., Jezequel, S., Meloux, G., Flodrops, Y., Landrieaux, J., Bouthier, A., Thomas, S., Solan, B.D., Gouache, D., 2015. Platform development for drought tolerance evaluation of wheat in france. Procedia Environmental Sciences 29, 93–94. https://doi.org/10.1016/j.proenv.2015.07.176

Duru, M., Therond, O., Martin, G., Martin-Clouaire, R., Magne, M.-A., Justes, E., Journet, E.-P., Aubertot, J.-N., Savary, S., Bergez, J.-E., others, 2015. How to implement biodiversity-based agriculture to enhance ecosystem services: A review. Agronomy for Sustainable Development 35, 1259–1281. https://doi.org/10.1007/s13593-015-0306-1

Foley, J.A., DeFries, R., Asner, G.P., Barford, C., Bonan, G., Carpenter, S.R., Chapin, F.S., Coe, M.T., Daily, G.C., Gibbs, H.K., others, 2005. Global consequences of land use. Science 309, 570–574. https://doi.org/10.1126/science.1111772

Gélard, W., Devy, M., Herbulot, A., Burger, P., 2017. Model-based Segmentation of 3D Point Clouds for Phenotyping Sunflower Plants, in: 12th International Joint Conference on Computer Vision, Imaging and Computer Graphics Theory and Applications (VISAPP 2017). Porto, Portugal, pp. 459–467. https://doi.org/10.5220/0006126404590467

Givry, S. de, Bouchez, M., Chabrier, P., Milan, D., Schiex, T., 2004. CARHTA GENE: Multipopulation integrated genetic and radiation hybrid mapping. Bioinformatics 21, 1703–1704. https://doi.org/10.1093/bioinformatics/bti222

Granier, C., Aguirrezabal, L., Chenu, K., Cookson, S.J., Dauzat, M., Hamard, P., Thioux, J.-J., Rolland, G., Bouchier-Combaud, S., Lebaudy, A., Muller, B., Simonneau, T., Tardieu, F., 2005. PHENOPSIS, an automated platform for reproducible phenotyping of plant responses to soil water deficit inArabidopsis thalianapermitted the identification of an accession with low sensitivity to soil water deficit. New Phytologist 169, 623–635. https://doi.org/10.1111/j.1469-8137.2005.01609.x

Hammer, G., Cooper, M., Tardieu, F., Welch, S., Walsh, B., Eeuwijk, F. van, Chapman, S., Podlich, D., 2006. Models for navigating biological complexity in breeding improved crop plants. Trends in Plant Science 11, 587–593. https://doi.org/10.1016/j.tplants.2006.10.006

Heslot, N., Akdemir, D., Sorrells, M.E., Jannink, J.-L., 2014. Integrating environmental covariates and crop modeling into the genomic selection framework to predict genotype by environment interactions. Theoretical and Applied Genetics 127, 463–480. https://doi.org/10.1007/s00122-013-2231-5

Hiederer, R., 2013. Mapping soil properties for europe: Spatial representation of soil database attributes. JRC, Luxembourg: Publications Office of the European Union, EUR26082EN Scientific; Technical Research series, ISSN 1831-9424; Citeseer. https://doi.org/10.2788/94128

Hiltunen, H.-M., Illarionov, B., Hedtke, B., Fischer, M., Grimm, B., 2012. Arabidopsis RIBA proteins: Two out of three isoforms have lost their bifunctional activity in riboflavin biosynthesis. International Journal of Molecular Sciences 13, 14086–14105. https://doi.org/10.3390/ijms131114086

Honsdorf, N., March, T.J., Berger, B., Tester, M., Pillen, K., 2014. High-throughput phenotyping to detect drought tolerance QTL in wild barley introgression lines. PLoS ONE 9, e97047. https://doi.org/10.1371/journal.pone.0097047

Hu, P., Chapman, S.C., Wang, X., Potgieter, A., Duan, T., Jordan, D., Guo, Y., Zheng, B., 2018. Estimation of plant height using a high throughput phenotyping platform based on unmanned aerial vehicle and self-calibration: Example for sorghum breeding. European Journal of Agronomy 95, 24–32. https://doi.org/10.1016/j.eja.2018.02.004

Husson, F., Lê, S., Pagès, J., 2010. Exploratory multivariate analysis by example using r. CRC Press. https://doi.org/10.1201/b10345

Jeuffroy, M.-H., Casadebaig, P., Debaeke, P., Loyce, C., Meynard, J.-M., 2013. Agronomic model uses to predict cultivar performance in various environments and cropping systems. A review. Agronomy for Sustainable Development 34, 121–137. https://doi.org/10.1007/s13593-013-0170-9

Lasky, J.R., Upadhyaya, H.D., Ramu, P., Deshpande, S., Hash, C.T., Bonnette, J., Juenger, T.E., Hyma, K., Acharya, C., Mitchell, S.E., Buckler, E.S., Brenton, Z., Kresovich, S., Morris, G.P., 2015. Genome-environment associations in sorghum landraces predict adaptive traits. Science Advances 1, e1400218–e1400218. https://doi.org/10.1126/sciadv.1400218

Lecoeur, J., Poiré-Lassus, R., Christophe, A., Pallas, B., Casadebaig, P., Debaeke, P., Vear, F., Guilioni, L., 2011. Quantifying physiological determinants of genetic variation for yield potential in sunflower. SUNFLO: a model-based analysis. Functional Plant Biology 38, 246–259. https://doi.org/10.1071/fp09189

Lemaire, G., Meynard, J.M., 1997. Use of the nitrogen nutrition index for the analysis of agronomical data, in: Lemaire, G. (Ed.), Diagnosis of the Nitrogen Status in Crops. Springer Berlin Heidelberg, Berlin, Heidelberg, pp. 45–55. https://doi.org/10.1007/978-3-642-60684-7_2

Lenz-Wiedemann, V., Klar, C., Schneider, K., 2010. Development and test of a crop growth model for application within a global change decision support system. Ecological Modelling 221, 314–329. https://doi.org/10.1016/j.ecolmodel.2009.10.014

Li, P., Li, Y.-j., Wang, B., Yu, H.-m., Li, Q., Hou, B.-k., 2016. TheArabidopsisUGT87A2, a stress-inducible family 1 glycosyltransferase, is involved in the plant adaptation to abiotic stresses. Physiologia Plantarum 159, 416–432. https://doi.org/10.1111/ppl.12520

Li, P., Li, Y.-J., Zhang, F.-J., Zhang, G.-Z., Jiang, X.-Y., Yu, H.-M., Hou, B.-K., 2016. The arabidopsis UDP-glycosyltransferases UGT79B2 and UGT79B3, contribute to cold, salt and drought stress tolerance via modulating anthocyanin accumulation. The Plant Journal 89, 85–103. https://doi.org/10.1111/tpj.13324

Li, Q., Yu, H.-M., Meng, X.-F., Lin, J.-S., Li, Y.-J., Hou, B.-K., 2017. Ectopic expression of glycosyltransferase UGT76E11 increases flavonoid accumulation and enhances abiotic stress tolerance in arabidopsis. Plant Biology 20, 10–19. https://doi.org/10.1111/plb.12627

Liang, Y., Ward, S., Li, P., Bennett, T., Leyser, O., 2016. SMAX1-LIKE7 signals from the nucleus to regulate shoot development in arabidopsis via partially EAR motif-independent mechanisms. The Plant Cell tpc.00286.2016. https://doi.org/10.1105/tpc.16.00286

Manavella, P.A., Arce, A.L., Dezar, C.A., Bitton, F., Renou, J.-P., Crespi, M., Chan, R.L., 2006. Crosstalk between ethylene and drought signalling pathways is mediated by the sunflower hahb-4 transcription factor. The Plant Journal 48, 125–137. https://doi.org/10.1111/j.1365-313x.2006.02865.x

Mangin, B., Casadebaig, P., Cadic, E., Blanchet, N., Boniface, M.-C., Carrère, S., Gouzy, J., Legrand, L., Mayjonade, B., Pouilly, N., André, T., Coque, M., Piquemal, J., Laporte, M., Vincourt, P., Muños, S., Langlade, N.B., 2017. Genetic control of plasticity of oil yield for combined abiotic stresses using a joint approach of crop modeling and genome-wide association. Plant, Cell and Environment. https://doi.org/10.1111/pce.12961

Marchand, G., 2014. Etude des réseaux de régulation impliqués dans la réponse au stress hydrique: Caractérisation, contrôle génétique et rôle au cours de l’évolution du tournesol cultivé, helianthus annuus (PhD thesis). Université de Toulouse.

Martre, P., He, J., Le Gouis, J., Semenov, M.A., 2015. In silico system analysis of physiological traits determining grain yield and protein concentration for wheat as influenced by climate and crop management. Journal of Experimental Botany. https://doi.org/10.1093/jxb/erv049

Messina, C., Boote, K., Loffler, C., Jones, J., Vallejos, C., 2006. Model-assisted genetic improvement of crops. Working with Dynamic Crop Models: Evaluation, Analysis, Parameterization, and Applications 309–335.

Messina, C.D., Podlich, D., Dong, Z., Samples, M., Cooper, M., 2011. Yield–trait performance landscapes: From theory to application in breeding maize for drought tolerance. Journal of Experimental Botany 62, 855–868.

Mittler, R., 2006. Abiotic stress, the field environment and stress combination. Trends in Plant Science 11, 15–19. https://doi.org/10.1016/j.tplants.2005.11.002

Moriondo, M., Bindi, M., Kundzewicz, Z.W., Szwed, M., Chorynski, A., Matczak, P., Radziejewski, M., McEvoy, D., Wreford, A., 2010. Impact and adaptation opportunities for european agriculture in response to climatic change and variability. Mitigation and Adaptation Strategies for Global Change 15, 657–679. https://doi.org/10.1007/s11027-010-9219-0

Noda, S., Yamaguchi, M., Tsurumaki, Y., Takahashi, Y., Nishikubo, N., Hattori, T., Demura, T., Suzuki, S., Umezawa, T., 2013. ATL54, a ubiquitin ligase gene related to secondary cell wall formation, is transcriptionally regulated by MYB46. Plant Biotechnology 30, 503–509. https://doi.org/10.5511/plantbiotechnology.13.0905b

Panagos, P., Van Liedekerke, M., Jones, A., Montanarella, L., 2012. European soil data centre: Response to european policy support and public data requirements. Land use policy 29, 329–338.

Pereyra-Irujo, G.A., Gasco, E.D., Peirone, L.S., Aguirrezábal, L.A.N., 2012. GlyPh: A low-cost platform for phenotyping plant growth and water use. Functional Plant Biology 39, 905. https://doi.org/10.1071/fp12052

Pereyra-Irujo, G.A., Velázquez, L., Lechner, L., Aguirrezábal, L.A., 2008. Genetic variability for leaf growth rate and duration under water deficit in sunflower: Analysis of responses at cell, organ, and plant level. Journal of experimental botany 59, 2221–2232. https://doi.org/10.1093/jxb/ern087

Picheny, V., Casadebaig, P., Trépos, R., Faivre, R., Da Silva, D., Vincourt, P., Costes, E., 2017. Using numerical plant models and phenotypic correlation space to design achievable ideotypes. Plant, Cell & Environment. https://doi.org/10.1111/pce.13001

Pieruschka, R., Poorter, H., 2012. Phenotyping plants: Genes, phenes and machines. Functional Plant Biology 39, 813. https://doi.org/10.1071/fpv39n11_in

Quesnel, G., Duboz, R., Ramat, É., 2009. The Virtual Laboratory Environment – An operational framework for multi-modelling, simulation and analysis of complex dynamical systems. Simulation Modelling Practice and Theory 17, 641–653. https://doi.org/10.1016/j.simpat.2008.11.003

Quilot-Turion, B., Génard, M., Valsesia, P., Memmah, M.-M., 2016. Optimization of allelic combinations controlling parameters of a peach quality model. Frontiers in Plant Science 7, 1873. https://doi.org/10.3389/fpls.2016.01873

Ramu, V.S., Paramanantham, A., Ramegowda, V., Mohan-Raju, B., Udayakumar, M., Senthil-Kumar, M., 2016. Transcriptome analysis of sunflower genotypes with contrasting oxidative stress tolerance reveals individual-and combined-biotic and abiotic stress tolerance mechanisms. PloS one 11, e0157522. https://doi.org/10.1371/journal.pone.0157522

Rasheed, A., Mujeeb-Kazi, A., Ogbonnaya, F.C., He, Z., Rajaram, S., 2017. Wheat genetic resources in the post-genomics era: Promise and challenges. Annals of Botany 121, 603–616. https://doi.org/10.1093/aob/mcx148

R Core Team, 2017. R: A language and environment for statistical computing. R Foundation for Statistical Computing, Vienna, Austria.

Rengel, D., Arribat, S., Maury, P., Martin-Magniette, M., Hourlier, T., Laporte, M., Varès, D., Carrère, S., Grieu, P., Balzergue, S., others, 2012. A gene-phenotype network based on genetic variability for drought responses reveals key physiological processes in controlled and natural environments. PloS one 7, e45249.

Robinson, M.D., McCarthy, D.J., Smyth, G.K., 2010. EdgeR: A bioconductor package for differential expression analysis of digital gene expression data. Bioinformatics 26, 139–140.

Sankaran, S., Khot, L.R., Espinoza, C.Z., Jarolmasjed, S., Sathuvalli, V.R., Vandemark, G.J., Miklas, P.N., Carter, A.H., Pumphrey, M.O., Knowles, N.R., Pavek, M.J., 2015. Low-altitude, high-resolution aerial imaging systems for row and field crop phenotyping: A review. European Journal of Agronomy 70, 112–123. https://doi.org/10.1016/j.eja.2015.07.004

Sarazin, V., Duclercq, J., Guillot, X., Sangwan, B., Sangwan, R.S., 2017. Water-stressed sunflower transcriptome analysis revealed important molecular markers involved in drought stress response and tolerance. Environmental and Experimental Botany 142, 45–53. https://doi.org/10.1016/j.envexpbot.2017.08.005

Segura, V., Vilhjálmsson, B.J., Platt, A., Korte, A., Seren, Ü., Long, Q., Nordborg, M., 2012. An efficient multi-locus mixed-model approach for genome-wide association studies in structured populations. Nature genetics 44, 825–830.

Sinclair, T., Hammer, G., Oosterom, E. van, 2005. Potential yield and water-use efficiency benefits in sorghum from limited maximum transpiration rate. Functional Plant Biology 32, 945–952.

Steduto, P., Hsiao, T.C., Raes, D., Fereres, E., 2009. AquaCropThe FAO crop model to simulate yield response to water: I. Concepts and underlying principles. Agronomy Journal 101, 426. https://doi.org/10.2134/agronj2008.0139s

Tardieu, F., 2011. Any trait or trait-related allele can confer drought tolerance: Just design the right drought scenario. Journal of Experimental Botany 63, 25–31. https://doi.org/10.1093/jxb/err269

Tardieu, F., Simonneau, T., Muller, B., 2018. The physiological basis of drought tolerance in crop plants: A scenario-dependent probabilistic approach. Annual review of plant biology.

Tilman, D., Cassman, K., Matson, P., Naylor, R., Polasky, S., 2002. Agricultural sustainability and intensive production practices. Nature 418, 671–677.

Underwood, J., Wendel, A., Schofield, B., McMurray, L., Kimber, R., 2017. Efficient in-field plant phenomics for row-crops with an autonomous ground vehicle. Journal of Field Robotics 34, 1061–1083. https://doi.org/10.1002/rob.21728

Van Ittersum, M., Rabbinge, R., 1997. Concepts in production ecology for analysis and quantification of agricultural input-output combinations. Field Crops Research 52, 197–208.

Velázquez, L., Alberdi, I., Paz, C., Aguirrezábal, L., Pereyra Irujo, G., 2017. Biomass allocation patterns are linked to genotypic differences in whole-plant transpiration efficiency in sunflower. Frontiers in plant science 8, 1976. https://doi.org/10.3389/fpls.2017.01976

Xu, Y., 2016. Envirotyping for deciphering environmental impacts on crop plants. Theoretical and Applied Genetics 129, 653–673. https://doi.org/10.1007/s00122-016-2691-5

Yu, J., Pressoir, G., Briggs, W.H., Bi, I.V., Yamasaki, M., Doebley, J.F., McMullen, M.D., Gaut, B.S., Nielsen, D.M., Holland, J.B., others, 2006. A unified mixed-model method for association mapping that accounts for multiple levels of relatedness. Nature genetics 38, 203–208.

